# The accumulation of lncRNAs in hybrid with DNA in patients with psoriasis reveals a decrease in the levels of RNase HII transcripts in the skin

**DOI:** 10.1101/2021.10.12.464167

**Authors:** Ecmel Mehmetbeyoglu, Leila Kianmehr, Murat Borlu, Zeynep Yilmaz, Seyma Basar Kılıc, Hassan Rajabi-Maham, Serpil Taheri, Minoo Rassoulzadegan

## Abstract

Long functional non-coding RNAs (lncRNAs) have been in the limelight in aging research because short telomeres are associated with higher levels of TERRA (Telomeric Repeat containing RNA). The genomic instability caused in Immune-mediated inflammatory diseases (IMID) especially in patients with psoriasis, which lead to short telomeres in psoriasis lesions, is a mechanism leading to cell aging. Research on the fraction of TERRA in hybrid with DNA offers avenues for new strategies. Skin samples were fractionated to obtain the RNA associated with DNA as a R-loop structure. TERRA analysis was performed by RT-qPCR and RNA-seq analysis. The higher amount of TERRA levels attached with each chromosome end was found with psoriasis patients. The increased levels of TERRA linked with telomeres correlate with the decrease in the *RNase-HII* transcript which means the unresolved DNA/RNA hybrids may ultimately facilitate the formation of skin lesions. LncRNAs have multiple molecular functions, including the regulation of heterochromatin, which controls genome stability and epigenome shaping and may be used as a trans-generational prognostic marker in patients with psoriasis.

## Introduction

Psoriasis is a chronic, recurrent inflammatory disease of the skin characterized by abnormal proliferation of keratinocytes, vascular hyperplasia, and infiltration of inflammatory cells into the dermis and epidermis^1^. It is generally accepted that the central pathogenesis of psoriasis is the dysfunction of T lymphocytes affected by complex interactions between genetic and environmental factors such as trauma, infections, stress, medications, smoking and alcohol consumption. The disease affects both men and women^2^. The uncertain etiology of psoriasis characterized by scaly erythematous plaques that can cause significant physical and psychological distress and affect approximately 125 million people worldwide^3,4,5^. The localization of the skin lesions is determined according to the traumatic triggers, the most frequent local lesions are on the knee, the elbow, and the scalp^6^. The abundance of reported familial cases suggests a possible genetic predisposition^7^.

Keratinocytes normally proliferate over a period of 40 days, but in patients with psoriasis their proliferation accelerates to only 4-8 days^1^. For many years, it has been debated whether dermal inflammation or epidermal proliferation is the cause of abnormal keratinocyte proliferation in the pathogenesis^8,9^. Immune-mediated inflammatory diseases (IMID) collectively describe a group of unrelated disorders that share common inflammatory pathways^10^, such as inflammatory bowel disease (IBD), psoriasis (PS), rheumatoid arthritis (RA), spondyloarthritis, and uveitis which belong to the group of age-related non-communicable diseases (NCDs) ^10,11,12,13^. Chronic inflammation, oxidative stress and alterations in telomere length (TL) have been shown to be involved in various age-related NCDs^14–17^.

Telomeres consist of the TTAGGG repeating units of DNA sequences associated with specific proteins called Shelterin complex at the ends of each chromosome^18,16,15^. Recently, a lncRNA TERRA has been reported with UUAGGG repeats which are transcribed directly from the end of each chromosome and hybridize to the leading strand and form a DNA/RNA R-loop^19,20^. In addition, the remaining single-stranded lagging 3′ end invades the double-stranded telomeric helix, forming a T-loop structure^21,22,20^. A specific reverse transcriptase complex (TERT) carrying its integral RNA template (TERC) adds telomeric repeats to the 3’ ends of chromosomes^23,24^. This enzyme, which is responsible for maintaining telomeric sequences is active at the early stage of development and in adult stem cells but mainly inactive in human somatic cells (except in lymphocytes)^25^. Its activation is a marker of rapid cell division in many cancer cells (85–90% of all human cancers)^26,27^. Altered TL has been associated with aging and aging-related diseases^28^. Telomere dysfunctions due to breaking or shortening during lifespan ultimately leads to cell senescence with chromosomal instability^29,20,30^.

The rate of telomere shortening may be accelerated by inflammation and oxidative stress *in vitro* and participate in the aging process^31^. Wu et al.^32,33^ demonstrated that in T lymphocytes in the blood of patients with psoriasis an increase in telomerase activity associated with a reduction in TL compared to healthy controls^24^ and it was also confirmed in Peripheral Blood Mononuclear Cells (PBMCs)^34^.

It is proposed that TERRA regulates the structure of telomeric chromatin^30,19^, but the mechanisms by which TERRA functions in disease development are poorly defined. It is reported that TERRA is preferentially recruited to short telomeres^30,17,22,14^ increasing the formation of hybrid DNA/RNA telomeric structures (R-loop) with the higher levels of TERRA triggering telomere fragility. In this study, we focused on the role of TERRA in maintaining genome and epigenome labeling. We propose a model where TERRA held together inside the ends of chromosomes imposes epigenetic changes of repetitive elements. We recently reported^35^ in normal cells that a fraction of transcribed TERRA is strongly maintained as a DNA/RNA hybrid in the telomeres from which they are derived. Here, we applied the same protocol to dissect and assess the levels of lncRNA molecule fractions, which are strongly attached to DNA in lesional (LA) and non-lesional (NL) skin and blood samples of psoriasis patients. Our results here reveal the detection of a general increased level of lncRNAs in R-loop with genome which correlates with decreased levels of Ribonuclease HII (*RNase HII*) transcripts in the skin of the psoriasis patients. In our study, particularly the lengths of telomeres and expression levels of TERRA molecules of different chromosomes were determined in skin biopsies and blood samples of healthy controls compared to patients with psoriasis, in LA and NL tissue samples. Overall, our results reveal the detection of an increased level of TERRA-mediated R-loop association with telomeres in psoriasis, providing a mechanism for telomeres length shortening during psoriasis progression.

## Results

Twenty patients with chronic plaque psoriasis were included in the study. Ten of these patients were women and ten were men. The healthy group consisted of 15 people, eight women (53.33%) and seven men (46.67%). Three sample groups of lesional (LA) and non-lesional (NL) psoriasis and healthy control were analyzed. The mean age of the control groups was 42.6 and 39.5 years, respectively. There is no statistically significant difference between the groups in terms of age and sex (p> 0.05). The mean Psoriasis Area and Severity Index (PASI) of the patient group was 15.6.

### Telomere length is significantly reduced in lesional tissue in psoriasis

Clear differences were observed in Figure 1 between psoriasis (PSO) lesional and PSO non-lesional skin samples in terms of telomere length (TL). TL was found to be significantly shorter in PSO lesional compared to PSO non-lesional tissue and the control group (p<0.05). There was no significant difference in TL between PSO non-lesional skin samples from patients with psoriasis and healthy controls (Figure 1, Table 1).

**Figure 1:**
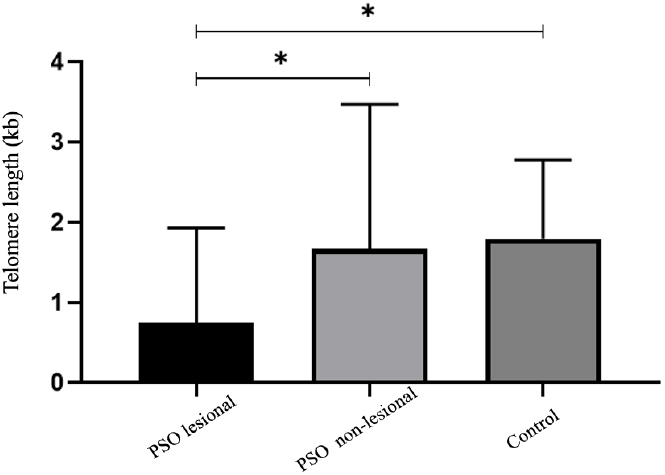
Telomere length (TL) in the skin. TL were determined by qPCR and TL of skin tissues in healthy control compared to PSO lesional and PSO non-lesional. Telomere length are significantly shorter in PSO lesional compared to PSO non-lesional(* p<0.05). Telomere length are significantly shorter in PSO lesional compared to health control (* p<0.05) PSO:Psoriasis, kb:kilobases

Persistent DNA damage with disease progression can lead to gradual changes in chromatin structure and erosion of the epigenetic landscape, which can be particularly harmful in regions of constitutive heterochromatin. According to several reports^17,23,28,29,31,34^ this potentially involves epigenetic alterations associated with aging in particular in the repetitive DNA sequences, including telomeric regions. We then tested the expression levels of lncRNA such as TERRA in the possible epigenetic alteration of TL.

### A higher level of TERRA is associated with telomeres in psoriasis patients

We designed experimental strategies to determine whether short telomere in LA tissue in psoriasis is associated with altered TERRA levels.

Recently, we described a simplified method of detecting DNA/RNA-hybrid regions in a complex genome. Unlike the often used DRIP techniques, it is not based on the recognition of the antigen or RNase H of the hybrid, but on a highly reproducible procedure to detect stable DNA/RNA hybrids that are maintained during the extraction protocol with TRIzol chloroform^35,36^. After extraction with TRIzol-chloroform the nucleic acids, are purified into two RNAs fractions, a small but reproducible amount of RNA is retained, along with high molecular weight genomic DNA, at the chloroform-water interface, while -free RNA molecules are found in the upper aqueous phase. The RNAs retained in DNA-RNA complexes were then recovered by chromatography on a “Zymo-SpinTM” column binding the DNA (Zymo-Research Corp Irvine CA, USA) followed by treatment with DNase. The routine free RNAs fraction was also purified from the aqueous phase after precipitation with isopropanol and further purification see Methods. Along with RT-qPCR, we examined the level of TERRA expression specifically transcribed from each chromosomal end on both fractions.

The aqueous phase extracted total RNAs (free fraction) from, both LA and NL skin samples expressed TERRAs (chromosomes 1q, 2q, 7p, 9p, 10q, 13q, 15q, 17p, 18p, X_q_Y_q_ and X_p_Y_p_ and total TERRA see Figure 2). The relative levels of TERRA expression showed low or no significant difference between the three groups of total free RNAs fractions, but in contrast, the DNA-bound fraction of TERRA profiles was significantly altered in the PSO lesional and PSO non-lesional skin tissue of affected patients with psoriasis. The difference is more significant in PSO lesional (see Figures 3). The DNA-bound RNA-hybrid (DRNA) obtained from PSO lesional and PSO non-lesional skin biopsy samples, compared to the control group shows a significant difference (*P*<0.05) in the relative levels expression of TERRA attached to the chromosomes 1q, 2q, 7p, 9p, 10q, 13q, 15q, 17p, 18p, XqYq, ,XpYp and sub-telomeric regions (Figure 3).

**Figure 2.**
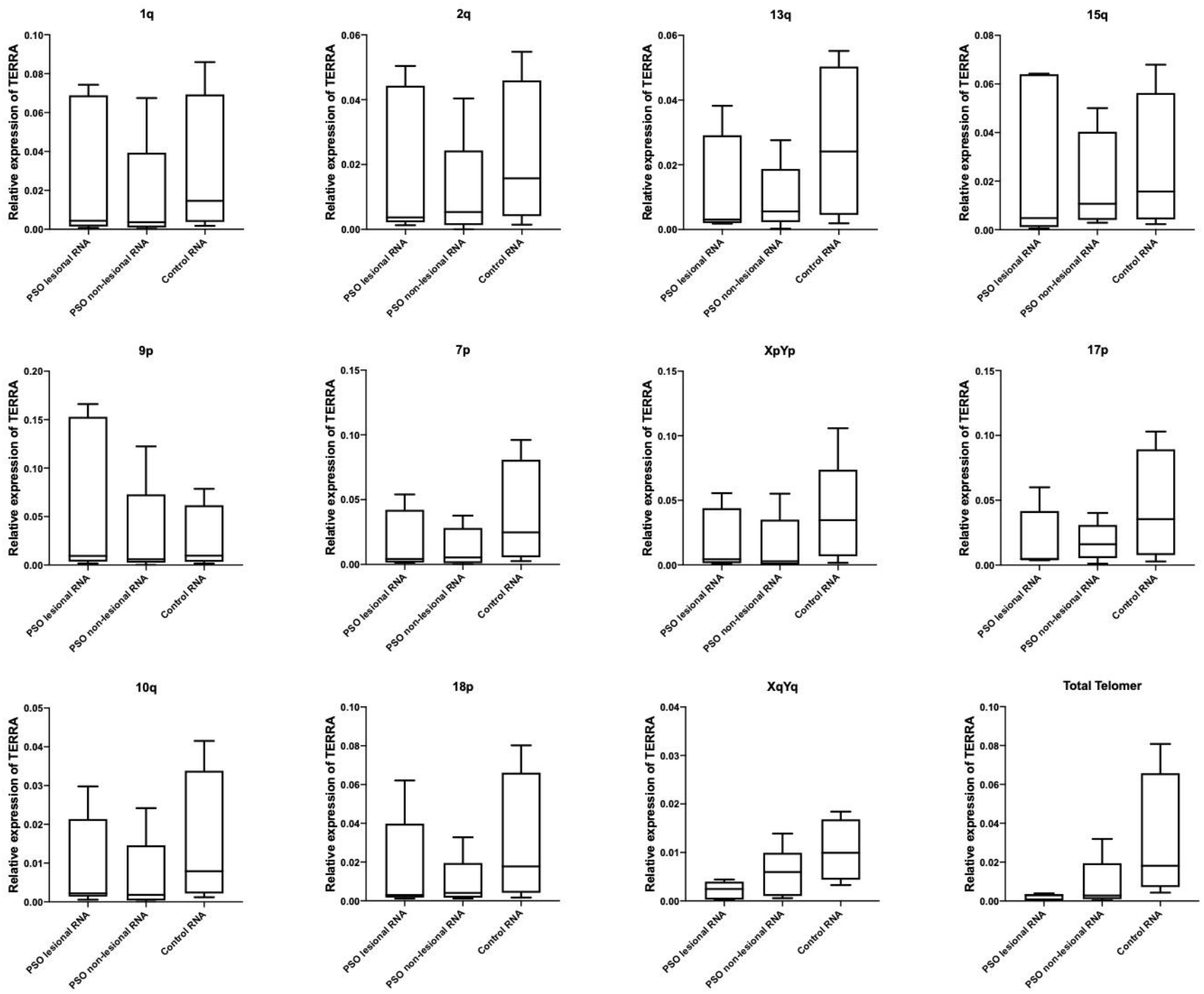
The levels of TERRA transcripts in aqueous phase (total free RNA). RT-qPCR analysis: were determined by relative normalized expression of TERRA in PSO lesional, PSO non-lesional and healthy control in terms of mRNA (total free-RNA fraction) of chromosomes: 1q, 2q, 7p, 9p, 10q, 13q, 15q, 17p, 18p, XqYq, XpYp, and Total TERRA. No difference in the levels of TERRA were found in psoriasis samples and control.

**Figure 3.**
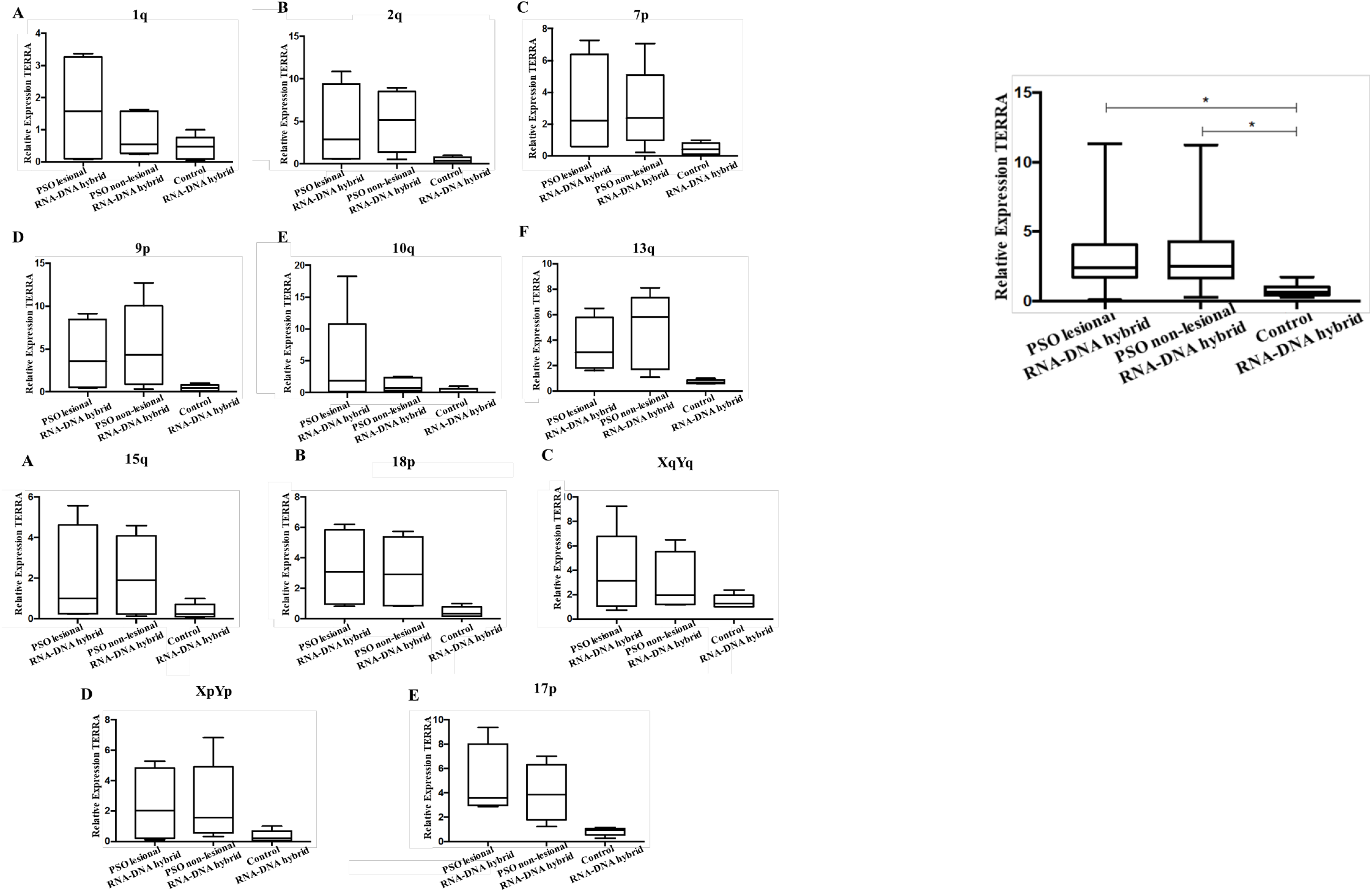
The levels of TERRA transcripts in DNA/RNA hybrid fraction. RT-qPCR analysis were determined by : relative normalized expression levels of TERRA in different chromosomes of psoriasis and healthy control DNA-RNA hybrids : 1q (A); 2q (B); 7p (C); 9p (D); 10q (E); 13q (F). : 15q (A); 18p (B); XqYq (C); XpYp (D); 17p (E). Significant differences in the levels of TERRA were found in psoriasis samples and control (* p<0.05).

Increases and decreases levels of TERRAs expression are associated with variable telomere length and can trigger uncontrolled cell division, genome instability, and cell senescence^14,37^.

Taken together, a higher level of TERRA was found in LA and NL DRNAs in psoriasis skin samples. This higher level is already detected in the non-lesional samples of patients with psoriasis, which means that the increase of TERRA in the fraction of RNA bound to DNA precedes the shortening of chromosomes length.

In addition, DNA and DRNA were isolated from both the patient’s blood and the healthy controls. Then, the length of the telomeres and the level of expression of TERRA were determined by RT-qPCR. According to our results, patients with PSO have a significantly shorter telomere length than the healthy controls. Contrary to these results, PSO patients have a significantly higher TERRA expression level. These results are consistent with the TL of the lesion tissue and the TERRA expressions results (see in Figure 4).

**Figure 4:**
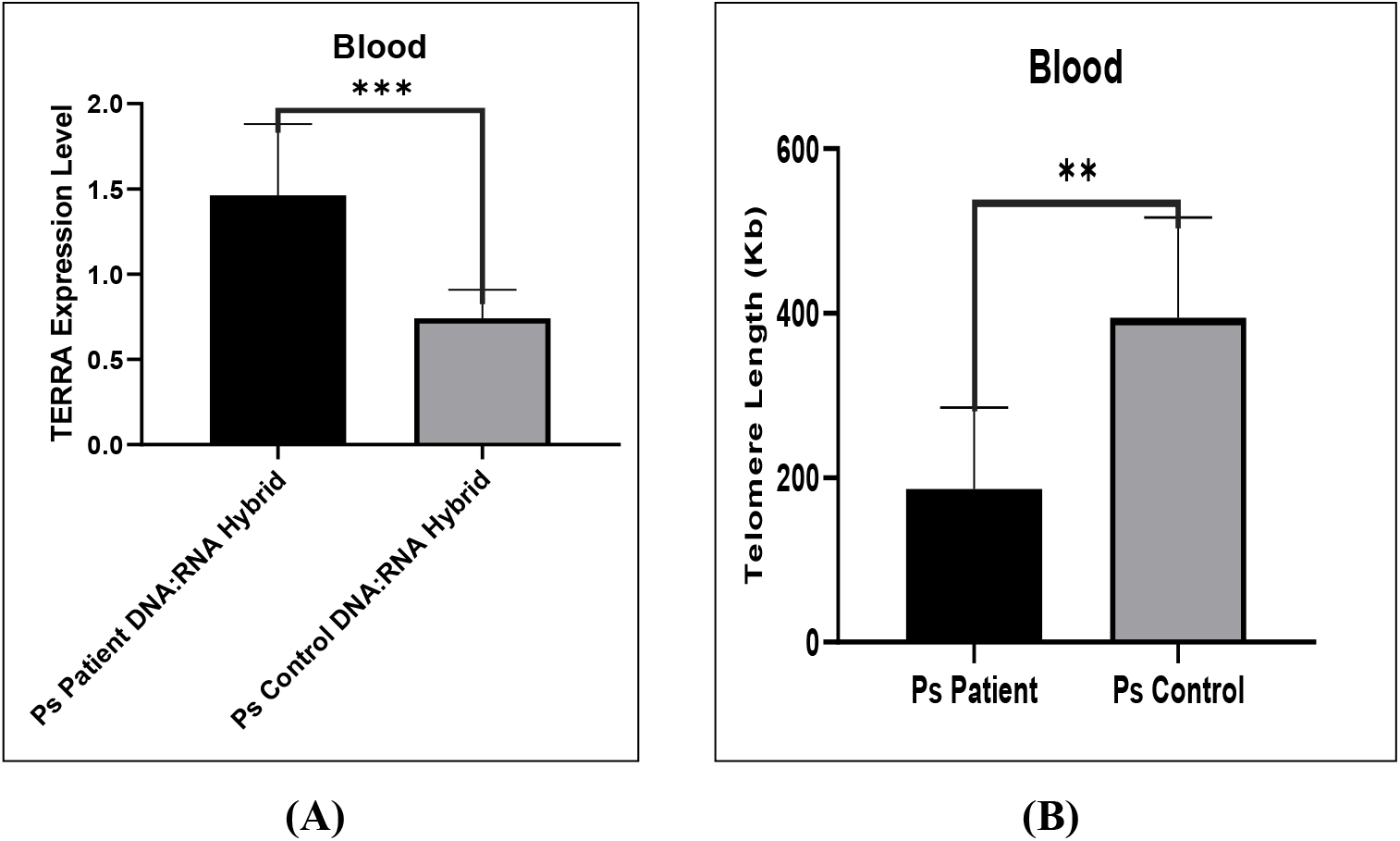
The levels of TL and TERRA transcripts in blood were determined by qPCR or RT-qPCR analysis: (A) The levels TERRA expression in DNA/RNA hybrid (p=0.0004) and (B) telomere length in blood tissue (p=0.061).

### Genome transcription profiles of patients with psoriasis: Analysis of the RNA fraction associated with DNA reveals higher levels of R-loop in the telomeric and centromeric regions

To get an overall view of LA and NL RNA profiling, RNA-seq analysis of DNA associated RNA fraction (DRNA) was performed. Recently, we reported that with a conventional sperm RNAs extraction protocol, unlike the free RNA fraction, a fraction made up of DNA/RNA hybrid molecules is tightly associated with the genome^35,36^. The DRNAs were recovered with the same way from skin samples of 6 healthy controls and 6 LA and NL samples of psoriasis patients.

To prepare the RNAs for the RNA-seq, the cutaneous cells were treated with proteinase K and then the DNA and RNAs fractions were recovered by column separation. The DNA retained on the columns was treated with DNase, and the RNA was released and further purified. The RNA fraction recovered from DNA is referred to as fraction D or DRNA for details see reference herein^35,36^. The DRNAs sequencing were performed by external service (see materials and methods). To identify significant peaks, we looked for genome-wide enriched peaks. The results are consistent between replicates, confirming that the technique is generally very reliable.

We assessed RNA sequences from human psoriasis skin cells of Lesions (LA), Normal (NL) and Control (C) individuals. LA and NL biopsies were taken from the same individuals. Interestingly, DRNA fraction from human somatic cells shows telomeric TERRA hybrids. Figure 5A-K shows RNA-seq signals over TERRA sequences at p- and q-arm telomeric regions of several human chromosomes. Each track shows a different sample (LA are DNA-bound RNA samples from lesions, NL are from normal and C are controls). The heights of the signals show scaled auto-group expression level over each genomic region using UCSC genome browser.

**Figure 5(A-K).**
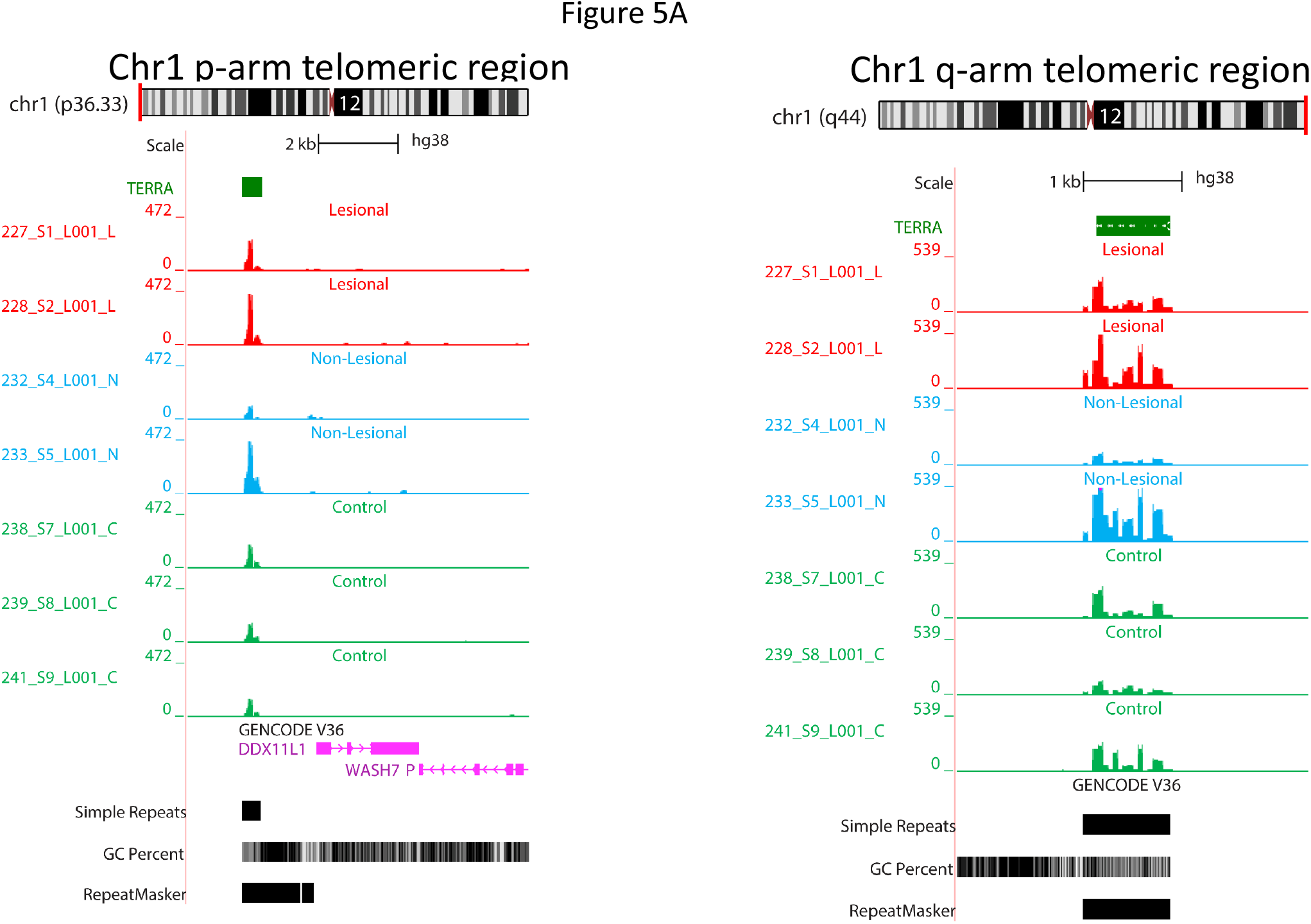
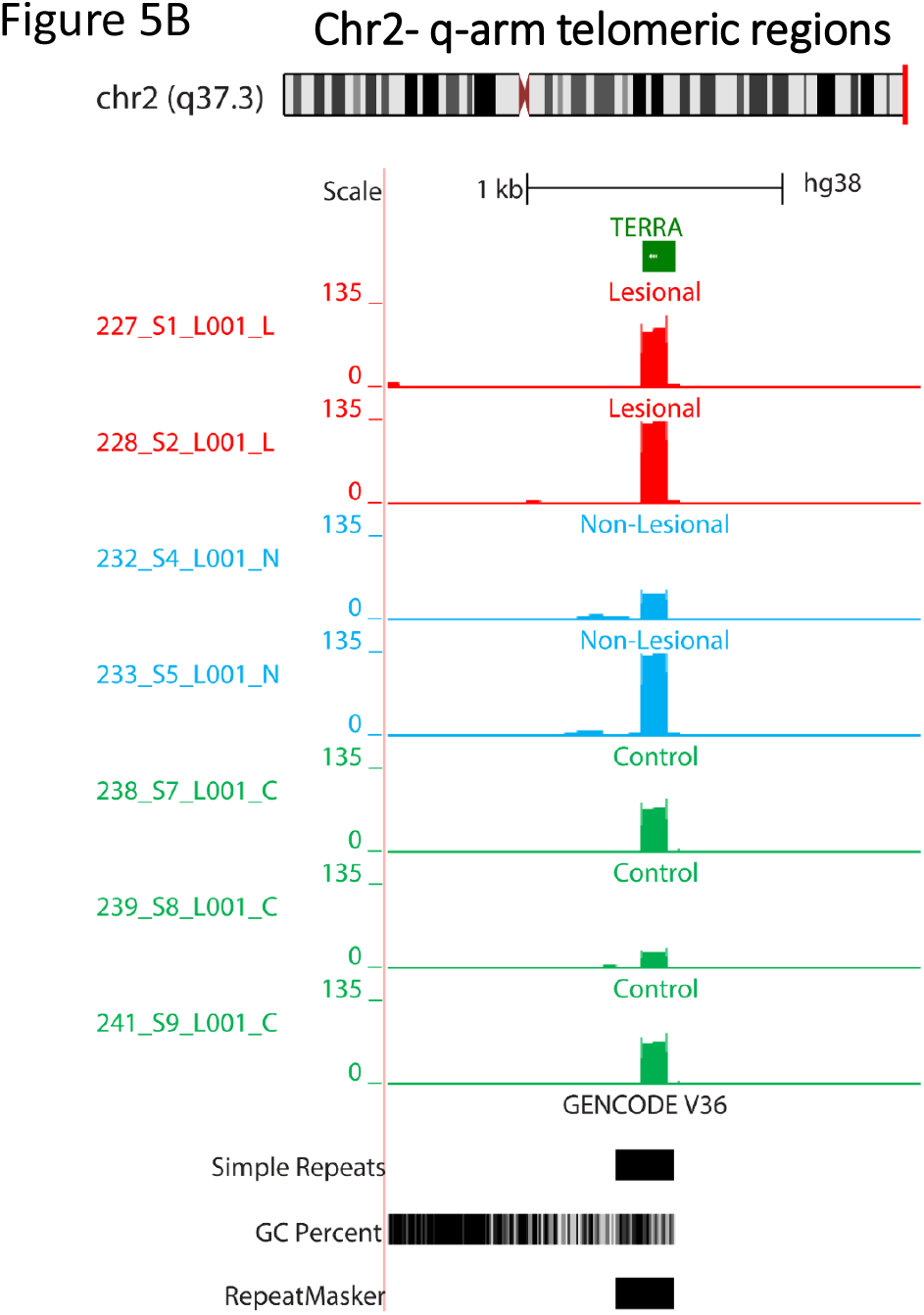
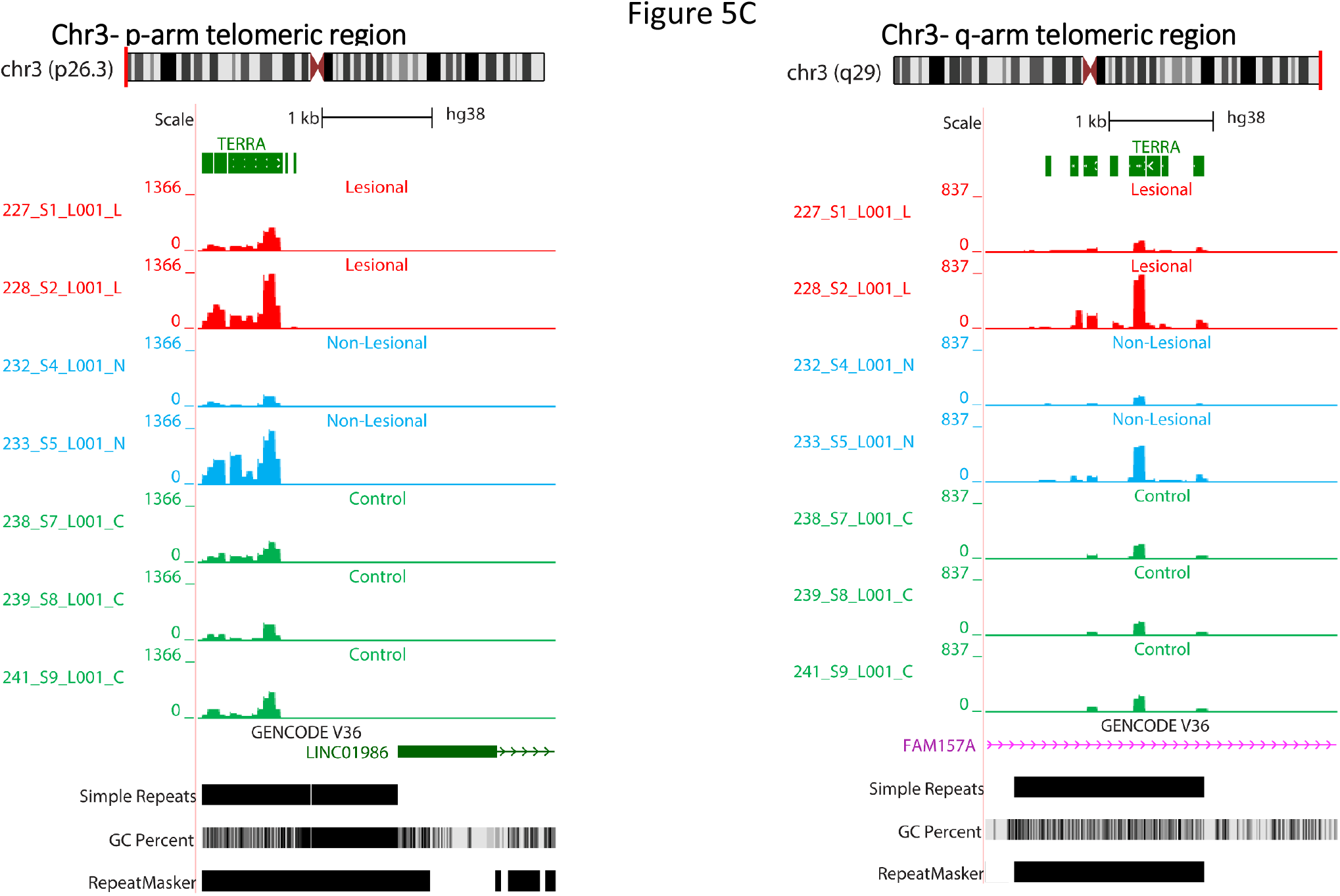
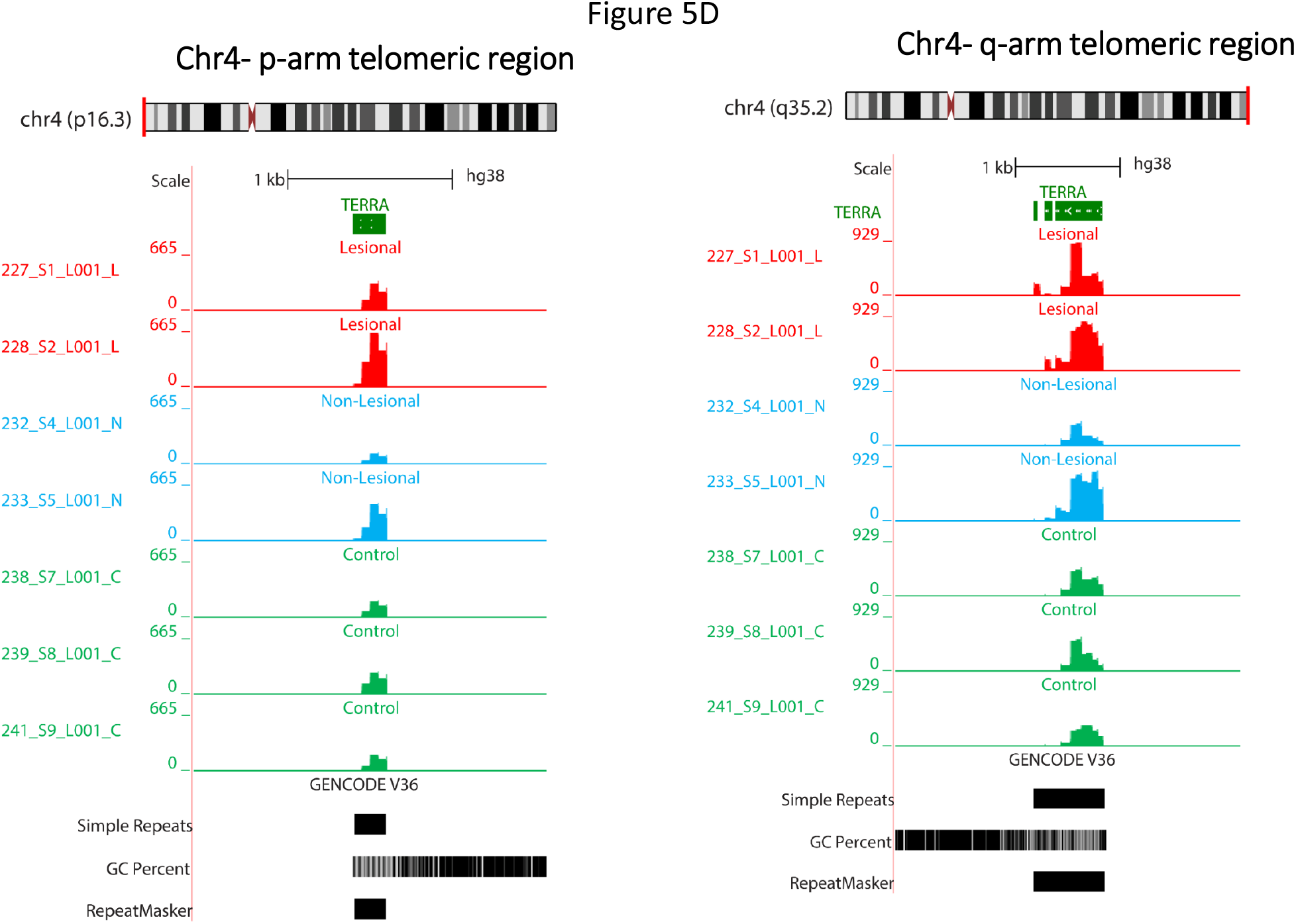
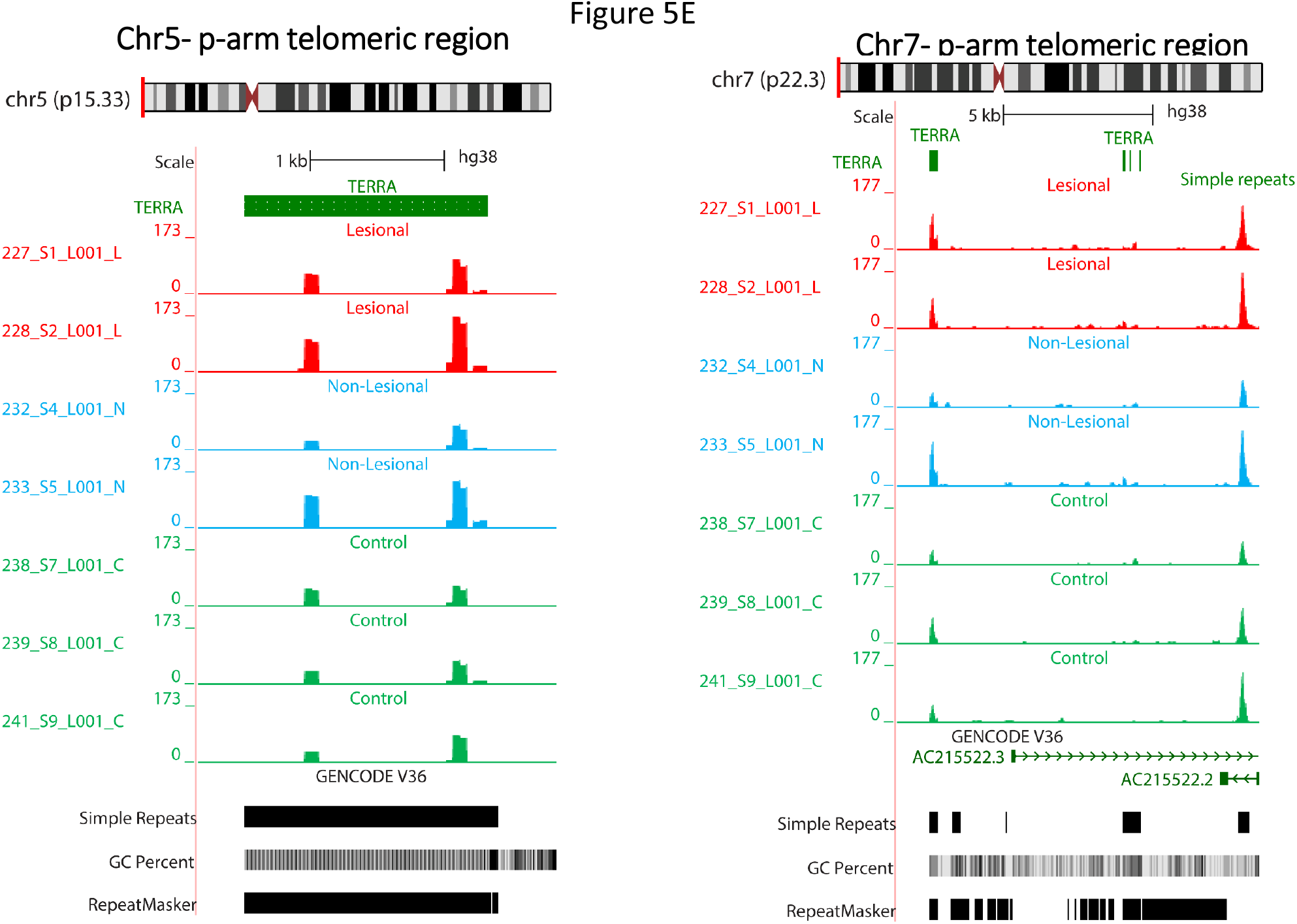
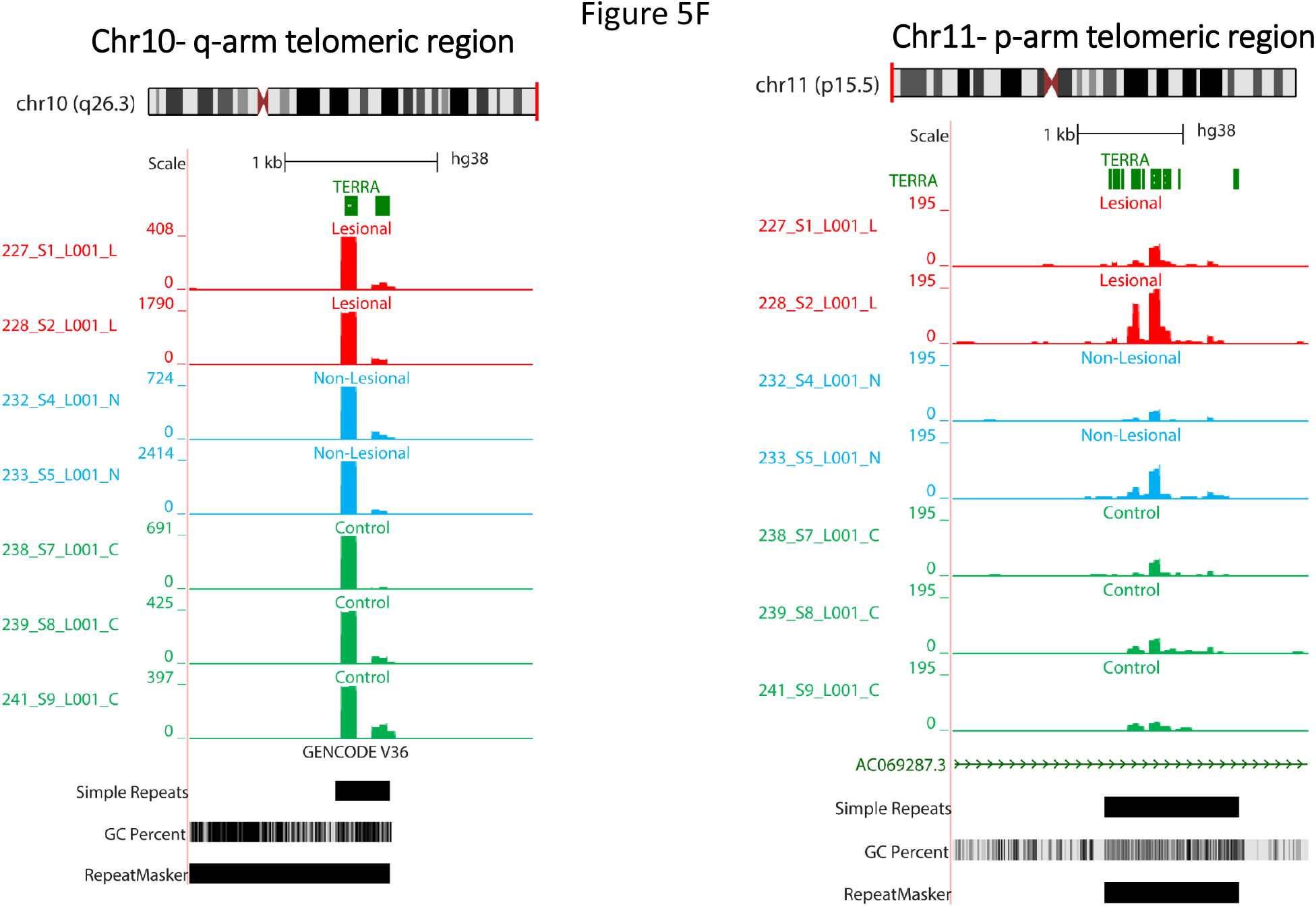
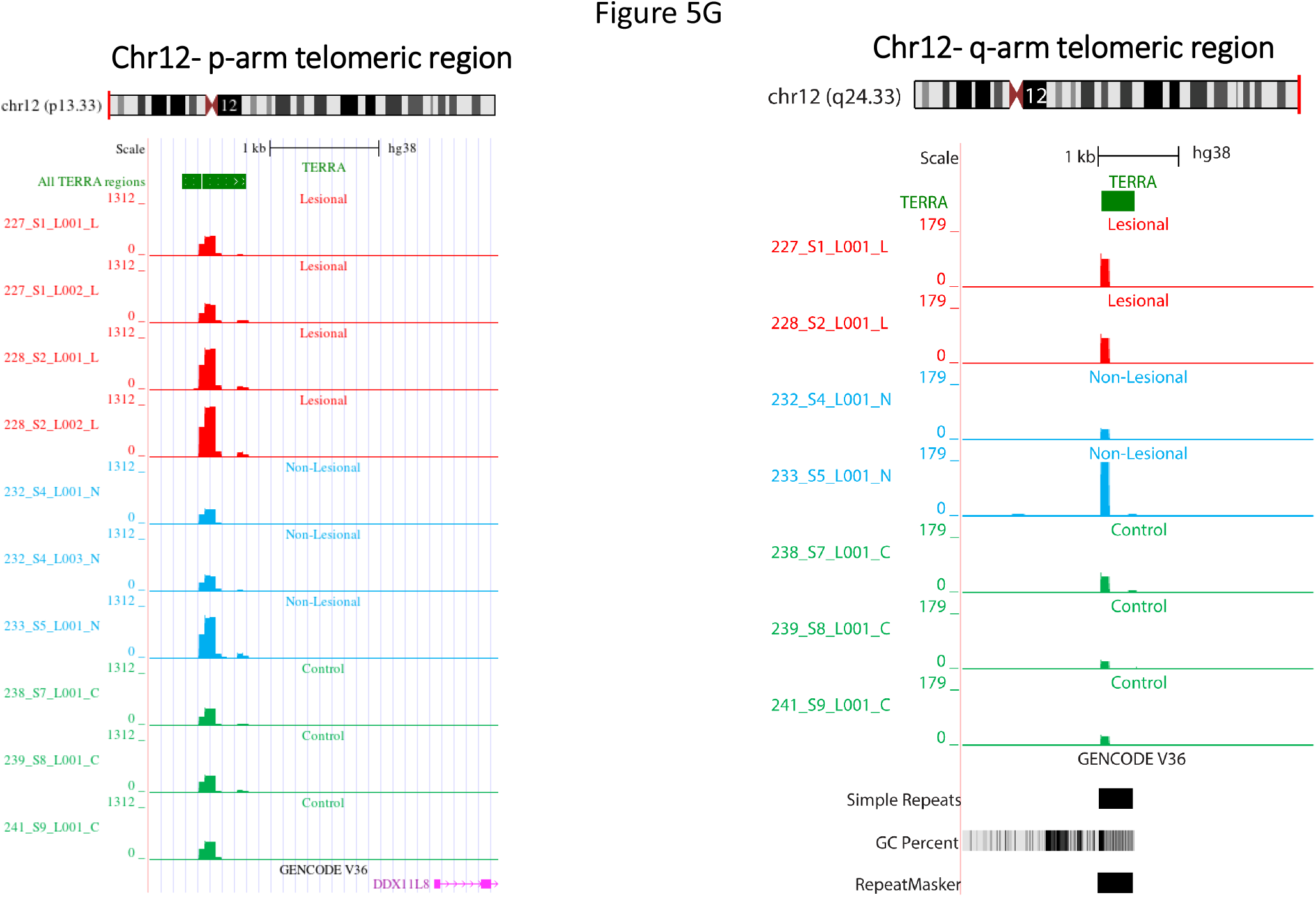
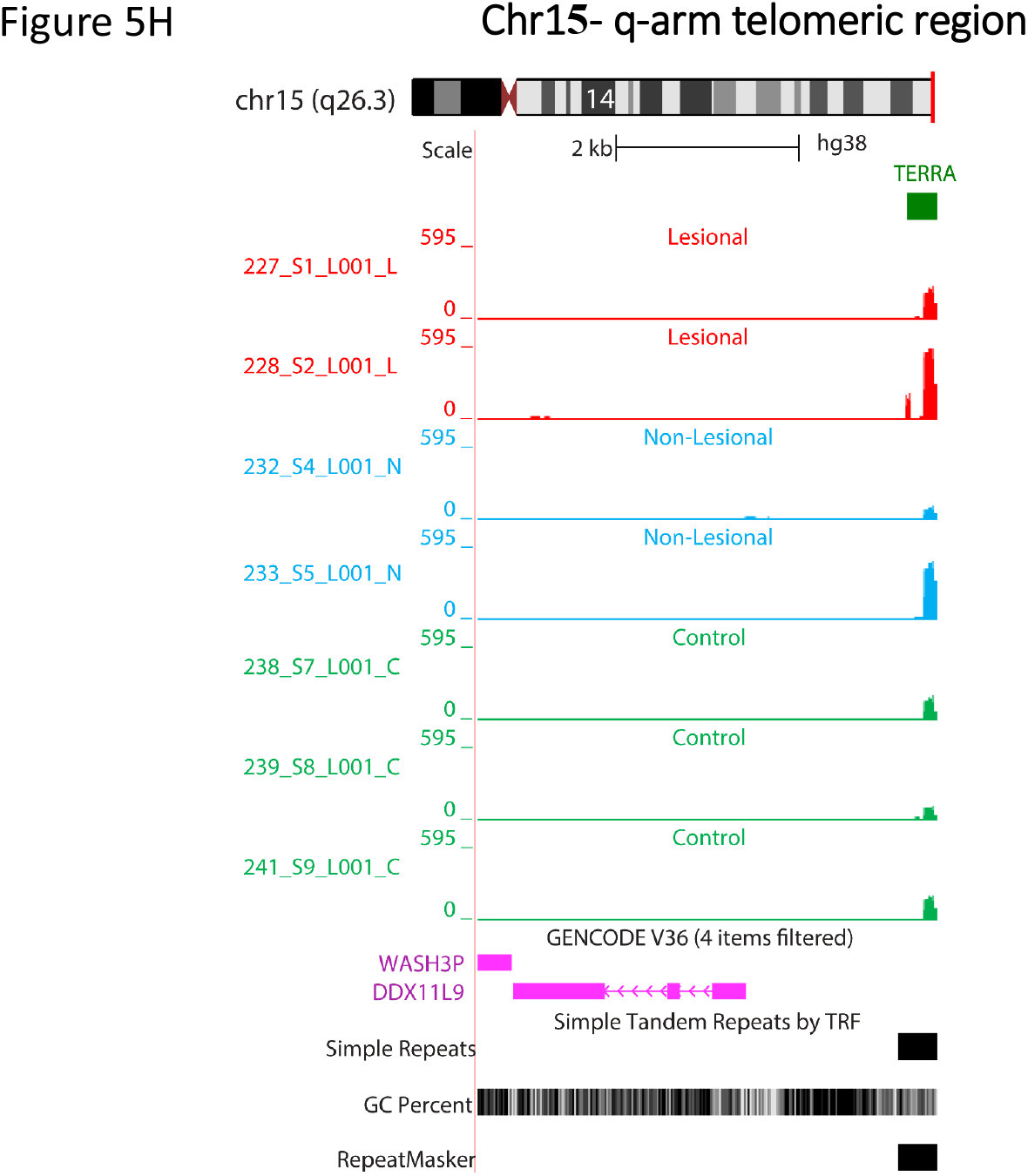
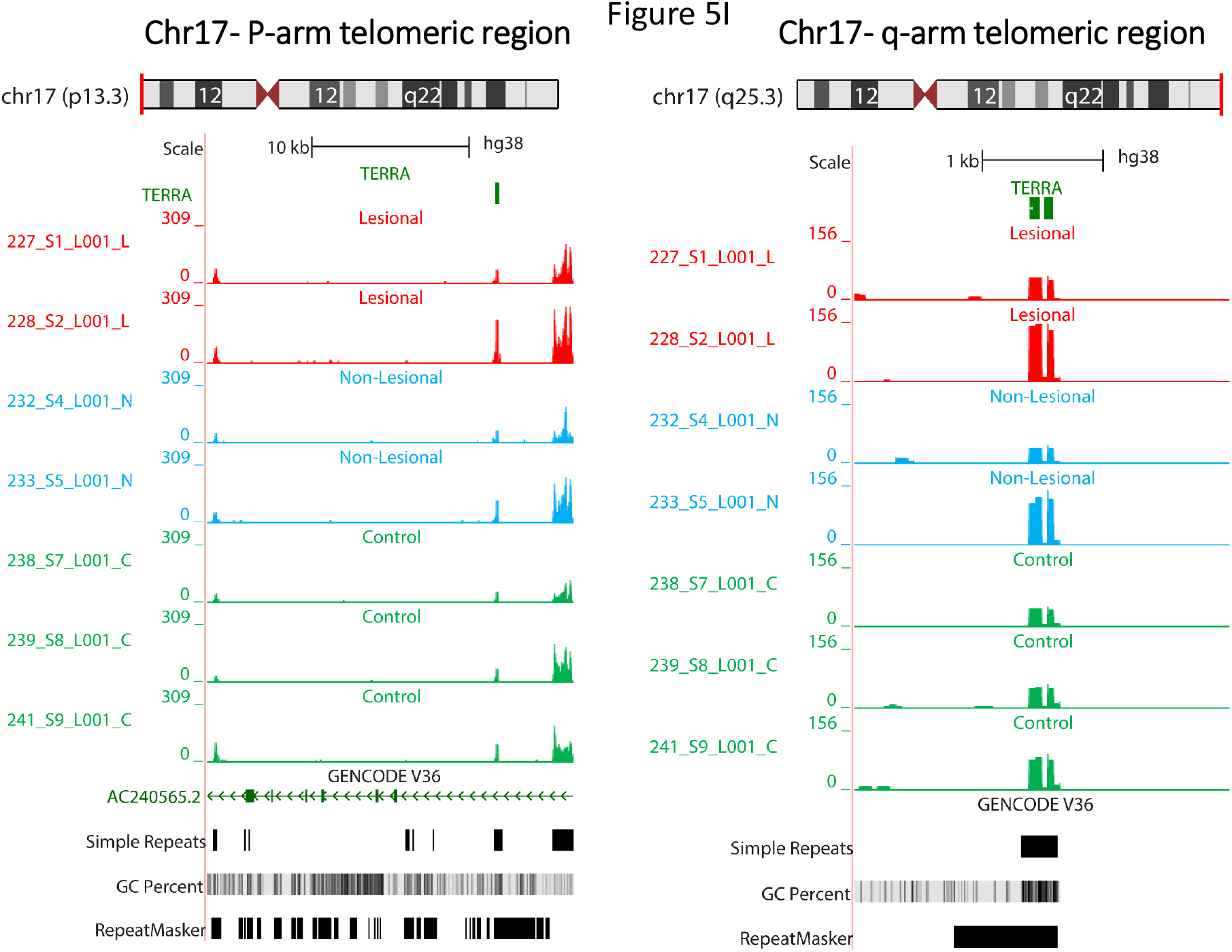
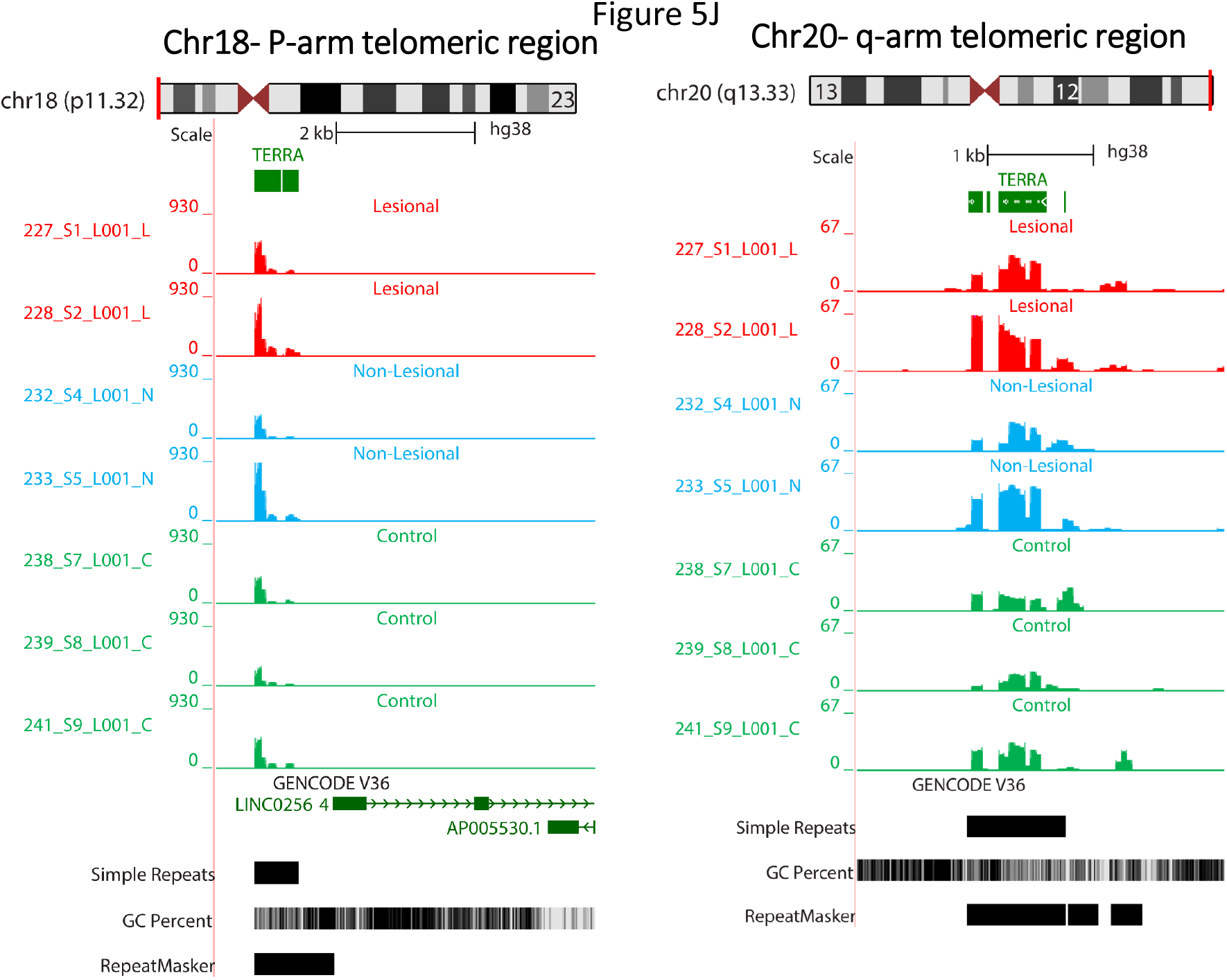
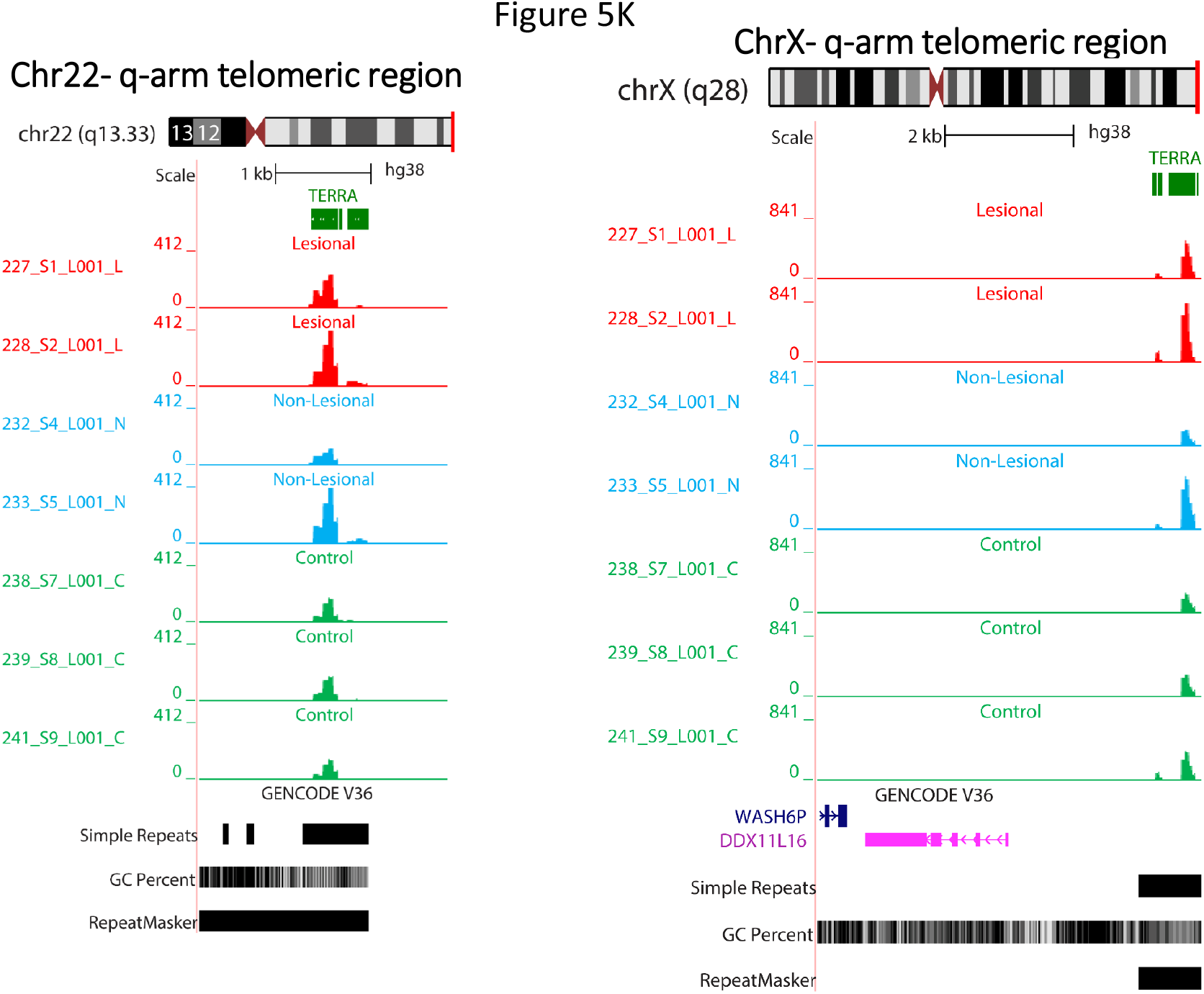
TERRA expression profiling genome-wide of DNA-bound RNA molecules over p-arm and q-arm chromosome telomeric regions. The Genome Browser figures show expression signal tracks of human DNA-bound RNA skin samples (Lesional(L), non-lesional(N), and controls(C)) over telomeric regions of p-arm and q-arm chromosomes (group auto-scaled view for all sample-group, each track is auto-scaled to display the track’s highest value). The upper track of each panel shows the chromosome number and the band which TERRA are located at, and the tiny green bar is the location of TERRA sequences in GRCh38 assembly (minimum four consecutive TAACCC repeats) of telomeric regions. The red, blue, and green tracks show expression signals in composite tracks (sample groups auto-scaled). The annotation track is also displayed in pack mode at the bottom of each window (Genecode V36). Coding exons are represented by blocks connected by horizontal lines representing introns. The 5’ and 3’ untranslated regions (UTRs) are displayed as thinner blocks on the leading and trailing ends of the aligning regions. In full display mode, arrowheads on the connecting intron lines indicating the orientation of the alignment pointing right if the query sequence was aligned to the forward strand of the genome and left if aligned to the reverse strand. Coloring for the gene annotation is based on the annotation types: coding, non-coding, Pseudogene, polyA annotations. Simple repeats, GC percent and repeat maskers are also shown at the bottom of all tracks.

#### 1-Telomeric regions

To view genome-wide TERRA loci throughout the different chromosomes, DNA/RNA hybrid density tracks along with TERRA loci (shown as a green bar) have been visualized on the ends of two arms of most of human chromosomes (Figure 5A-K). TERRA loci sequences (minimum four consecutive sequences TTACCC) loaded as a bed file with the BigWig data using the UCSC genome browser. These tracks display read density over telomeric TERRA regions based on hg38 genome assembly that reveal normalized TERRA expression signal. Each graph-based track is configured to highlight different samples (LA in red, NL in blue, and controls are in green colors). Read density tracks of all sample groups are auto-scaled to view normalized reads in comparison. All tracks are scaled against the one track that has the highest value (scales shown at the left of each browser). Signal tracks of LA, NL and controls revealed homogeneous expression signal at TERRA regions at p-arm or q-arm or both end of different human chromosomes (1p, 3p, 4p, 5p, 11p, 12p, 17p, 18p), and (1q, 2q, 3q, 4q,10q, 12q, 15q, 17q, 20q, 22q, Xq). TERRA signals are present at both ends of chromosomes 1, 3, 4, 12, 17. The comparative analysis of fully sequenced sub-telomeres of human chromosomes; 1p, 3p, 4p, 5p, 11p, 12p, 17p, 18p, 1q, 2q, 3q, 4q, 10q, 12q, 15q, 17q, 20q, 22q, Xq has revealed a common structure, in which the proximal and distal sub-telomeric domains are separated by TTAGGG repeats. TERRA repeats are indicated within upstream and downstream regions in average 1-50 kb on every chromosome ends to show a range of transcripts proximity.

RNA-seq expression profiling of LA, NL, and control samples shows that TERRA levels signals are relatively higher in LA and NL than from controls (Figure 5A-K). This observation also confirms the RT-qPCR results according to which the expression level of TERRA in the LA and NL samples groups is much higher than the control samples groups.

#### 2-TERRA transcripts in the individual telomeres from local sub-telomeric promoters of human somatic cells

Sub-telomeric regions of human chromosomes contain many transcripts, members of non-coding RNA and pseudogenes families. During the profiling of TERRA repeats, we found a family of pseudogene transcripts with DDX11L members which located in the sub-telomeric region (1-2 kb): DDX11L1 and WASH7P in 1p, DDX11L8 in 12p, DDX11L9 and WASH3P in 15q, DDX11L16 and WASH6P in Xq. The sequence alignment also revealed that DDX11L is overlapped to the WASH gene. The RNA-seq alignment also confirms the presence of these transcripts nearby telomeric regions (see figures 5A, 5G, 5H, and 5K). Costa et al also reported a novel gene family, indicating that an ancestral gene, originated as a rearranged portion of the primate DDX11 gene, and propagated along with many sub-telomeric locations of human chromosomes^38^. The presence of the gene family reveals sub-telomeres dynamics, and emergence of a multicopy transcript from an inactive pseudogene as in our data 3q TERRA repeats are overlapped to FAM157A that belongs to pseudogene biotype. In addition, Figure 5F, 5I show TERRA repeats in 11p and 17p overlapped to a non-coding RNA (AC069287.3 and AC240565.2 respectively).

Sub-telomeric transcripts of the DNA-bound RNA molecules may also occurs from every chromosome in human somatic cells as well as germ cells^35^.

#### 3-More lncRNA expression signals are observed in non telomeric regions of the lesional and non-lesional skin of patients with psoriasis

Moreover, we also reported in our previous study that in addition to telomeric TERRA regions there are some non-telomeric regions in sperm DRNA fractions^35^. Here they are not as much as the sperm DRNA fraction. For instance, non-telomeric TERRA region on chr2 can be observed that is overlapped to a long intergenic non-coding RNA (LINC01934) as shown in Figure 6.

**Figure 6.**
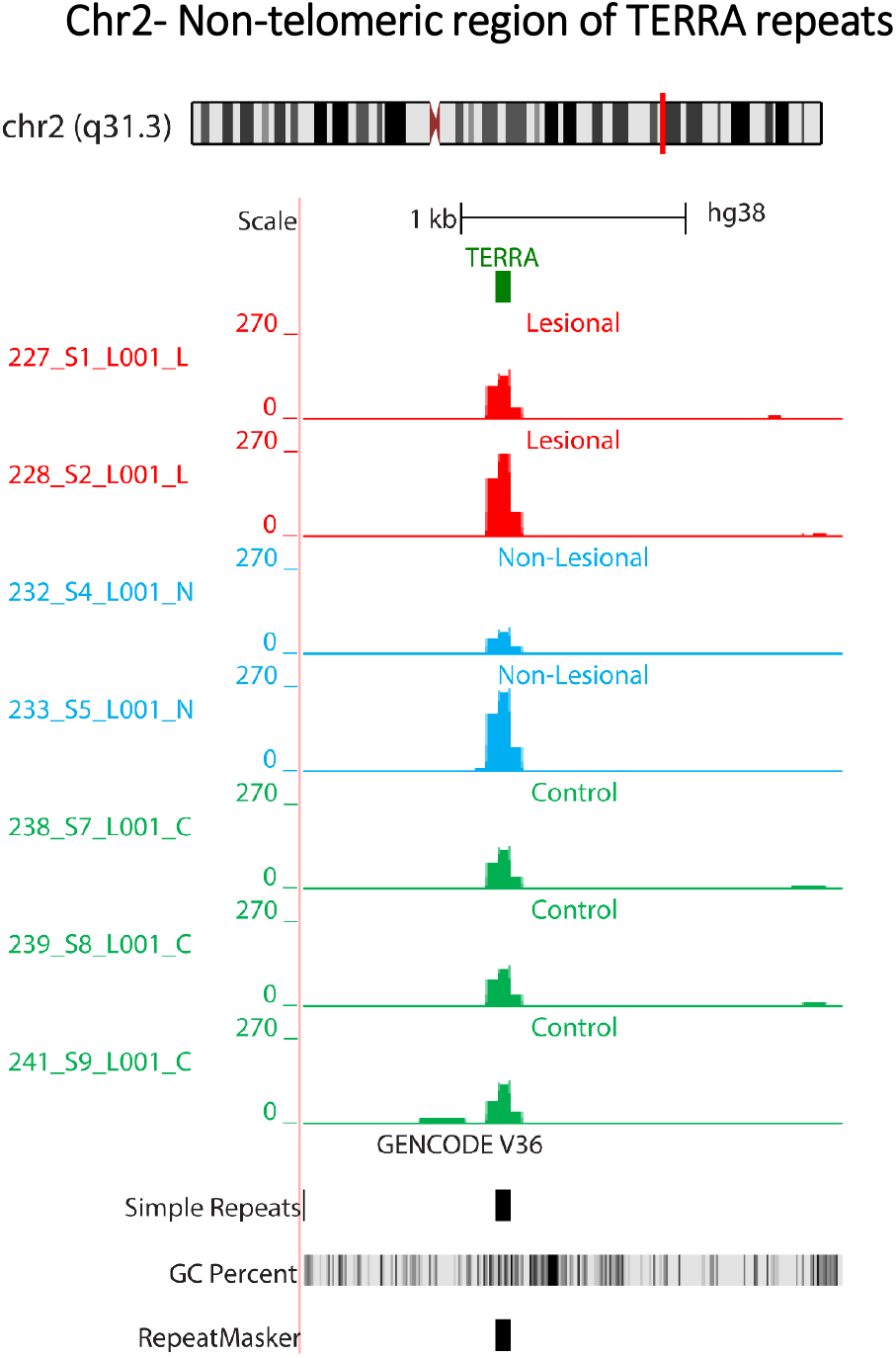
Non-telomeric region of TERRA sequences on chr2 (q31.3). The Genome Browser figures show read density tracks of human DNA-bound RNA skin samples (L, N, and C) over non-telomeric region of TERRA repeats on chromosome2 (group auto-scaled view for all sample-group, each track is auto-scaled to display the track’s highest value). The upper track of each panel shows the chromosome number and the band which TERRA are located at, and the tiny green bar is the location of TERRA sequences in GRCh38 genome assembly (minimum four consecutive TAACCC repeats) of telomeric regions. The red, blue, and green tracks show read density in composite tracks (sample groups auto-scaled). The annotation track is also displayed in pack mode at the bottom of each window (Genecode V36).

Annotated positions of exons, intergenic, intron, promoter-TSS (Transcription Start Site) and TTS (Transcription Termination Site) were based on the hg38 assembly genomic features which shows that most of the peaks in skin DRNA samples fall into enhancer regions (intergenic and intronic regions) compared to others. In conclusion, we have the same observation for skin DRNA as in human and mouse sperm data while most of the peaks in sperm free RNA and testes DRNA fall into the exonic regions^35^. Thus, it seems that we can see strong reproducibility in DRNA sperm (as a germ cell) and skin (as a somatic cell). The general transcripts of the regions: exon, promoter-TSS and TTS are perfectly comparable levels in all samples (see Figure 7).

**Figure 7.**
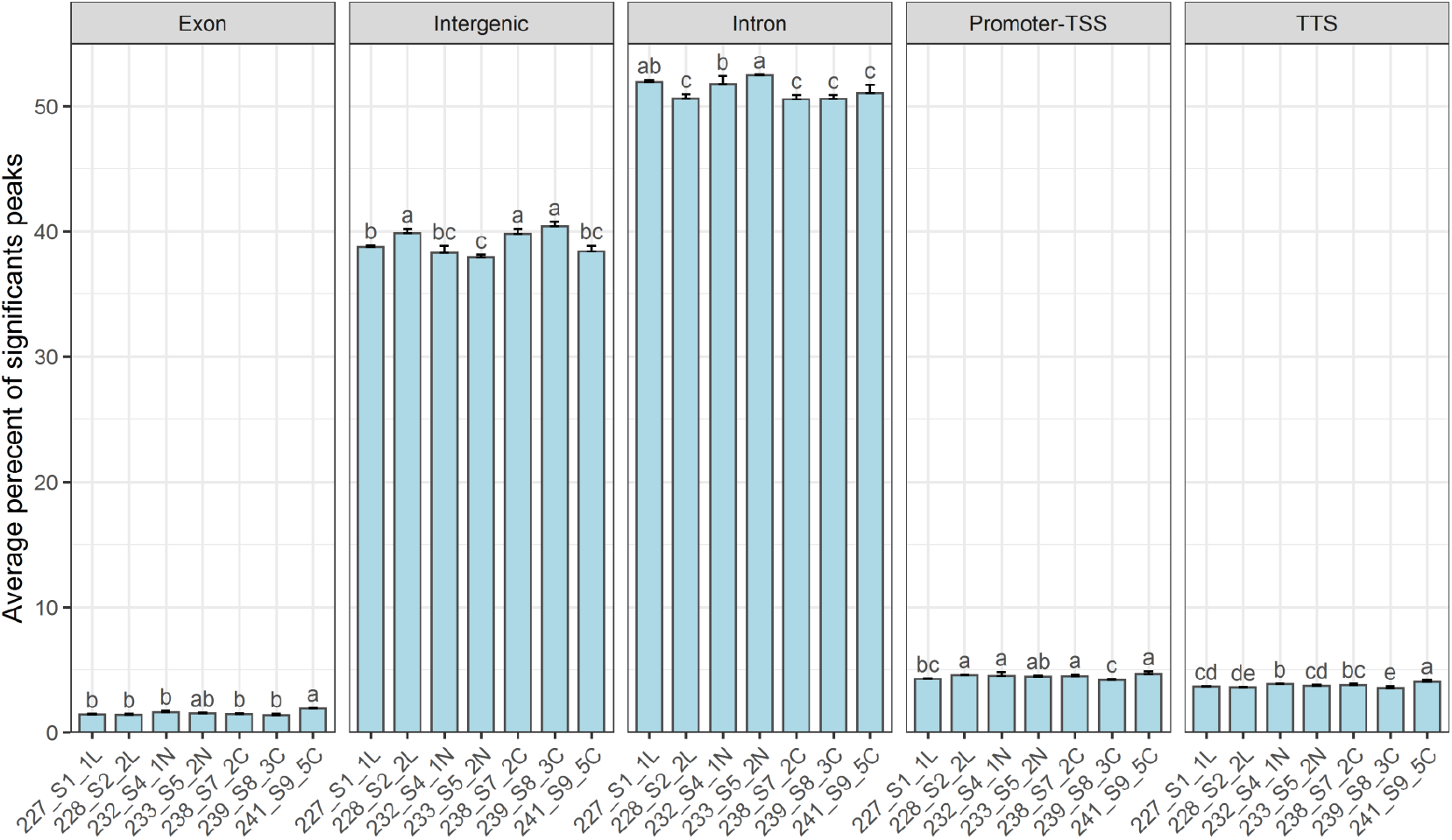
Peak finding. The Average percent of significant peaks of different genomic regions including exon, intergenic, intron, promoter-TSS, and TTS in all sample groups. Values are mean±SD of each sample group biological replicates. Bar plots with different superscript are significantly different (one-way ANOVA) at the level of p<0.05 followed by Duncan’s multiple comparison test. One-way ANOVA has been performed for each genomic region, respectively. (1L, 2L; lesional), (1N, 2N; non-lesional), (2C, 3C, 5C; healthy controls). Each sample group including 2-3 biological replicates with four technical replicates.

On the other hand, in accordance with the RT-qPCR technique results, higher expression values are also observed in non-lesional samples from psoriasis patients by the report to the control. But, the lesional samples profiling are more complex. More analysis is required to compare other genomic regions such as ribosomal etc. that can be considered for further studies.

#### 4-Transcripts from centromeric regions

Centromeric regions (see below) also accumulate more DNA-bound RNA compared to controls not only in lesion samples but also in the healthy samples from patients with psoriasis (see Figure 8A-D).

**Figure 8.**
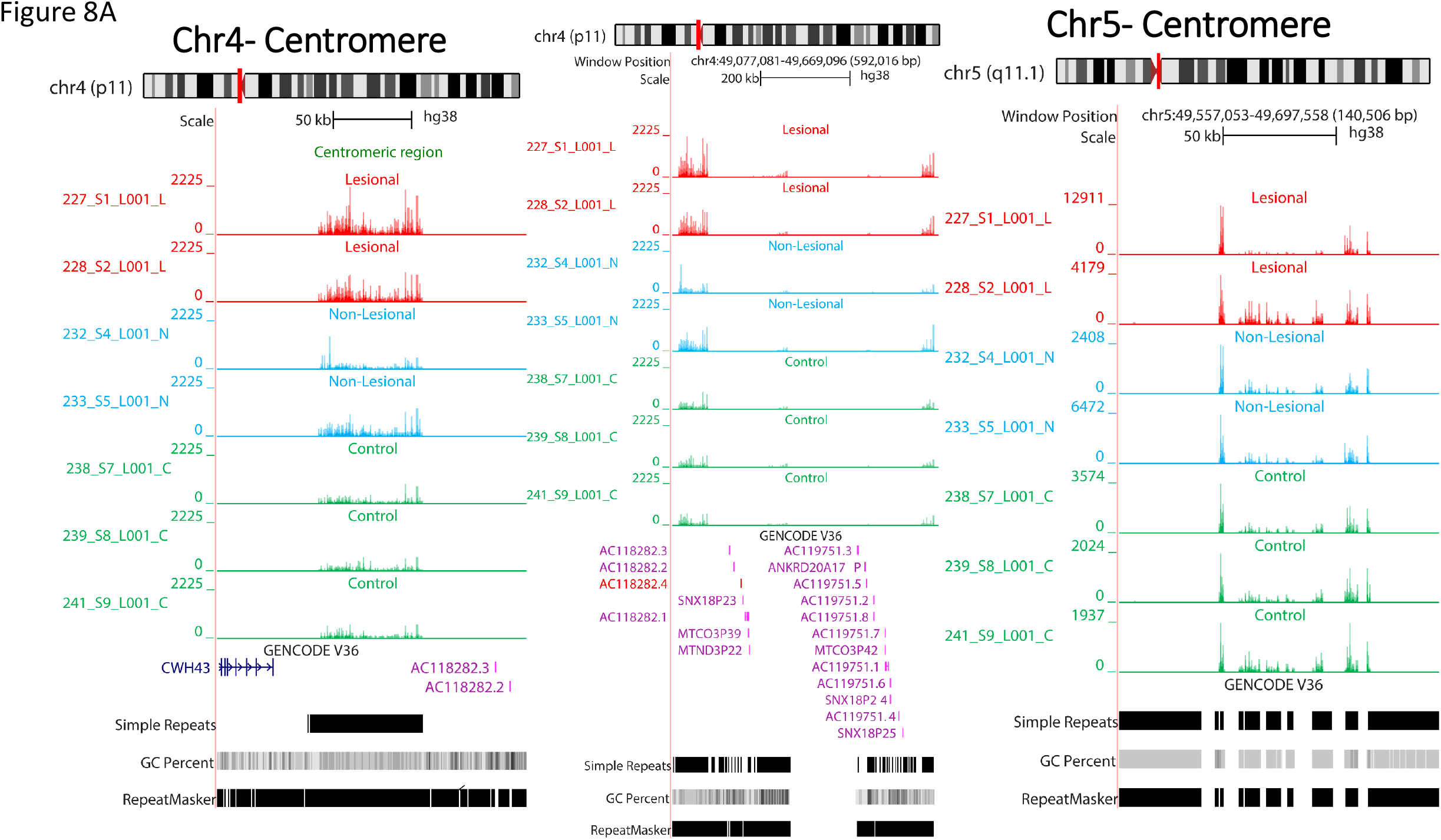
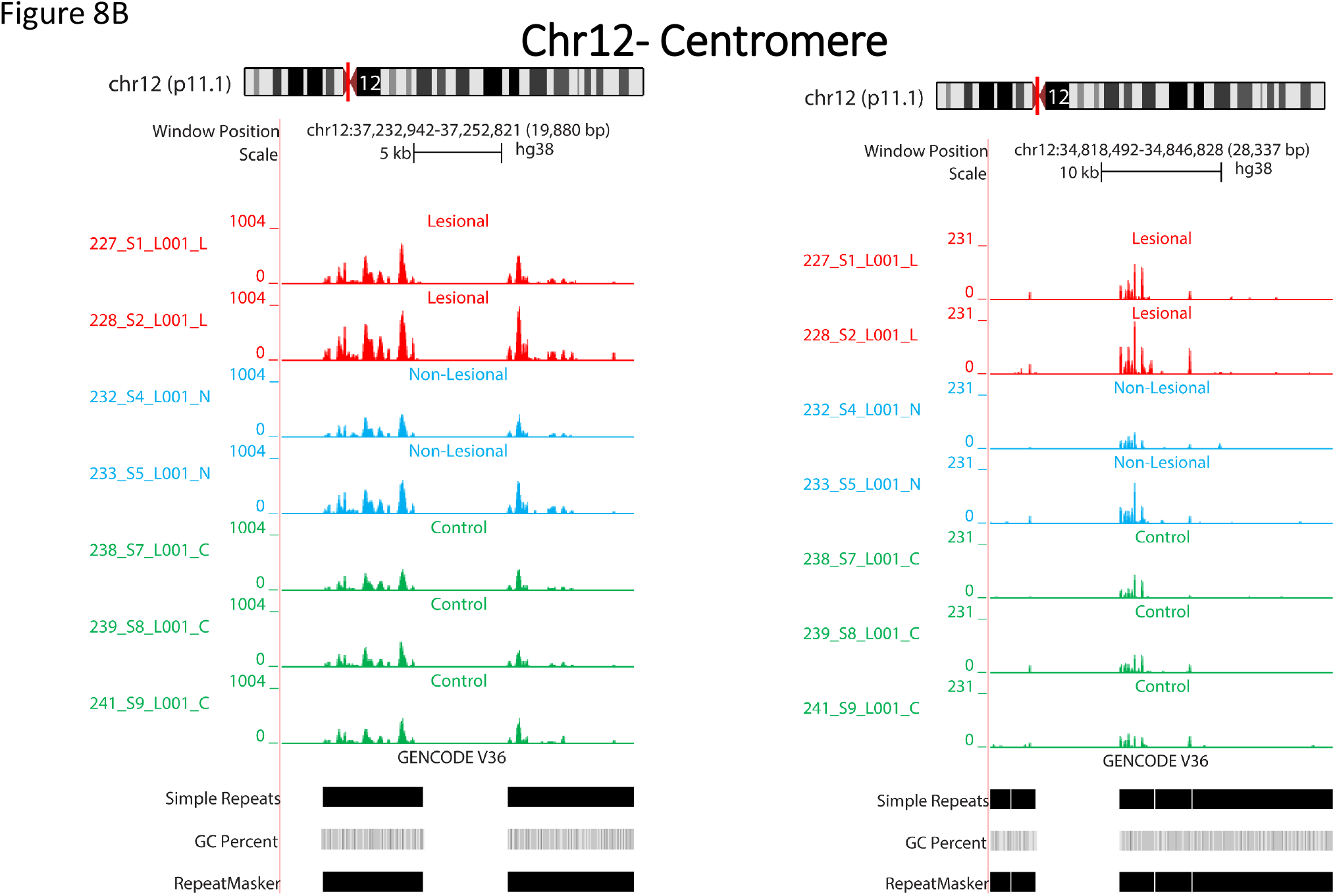
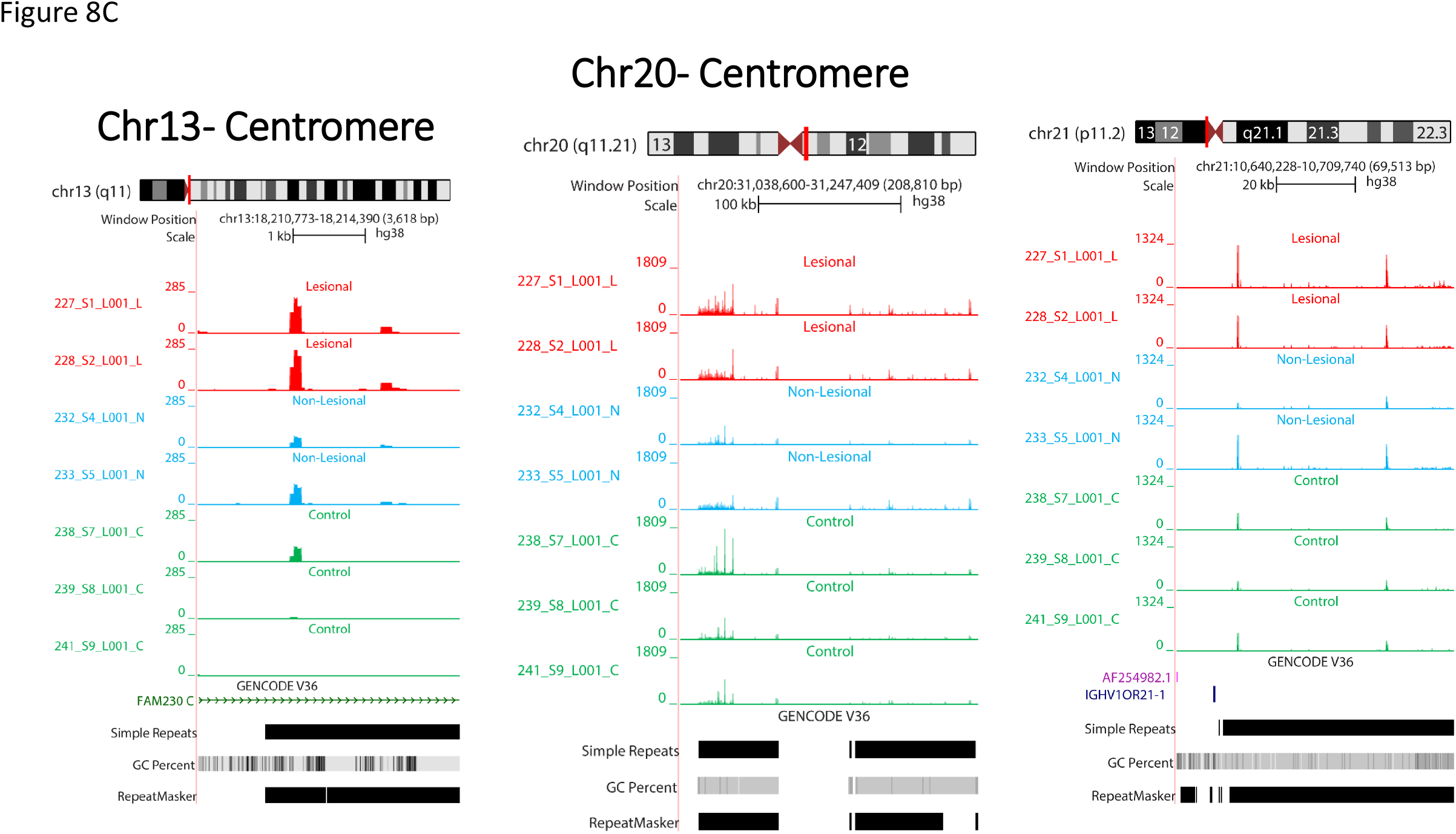
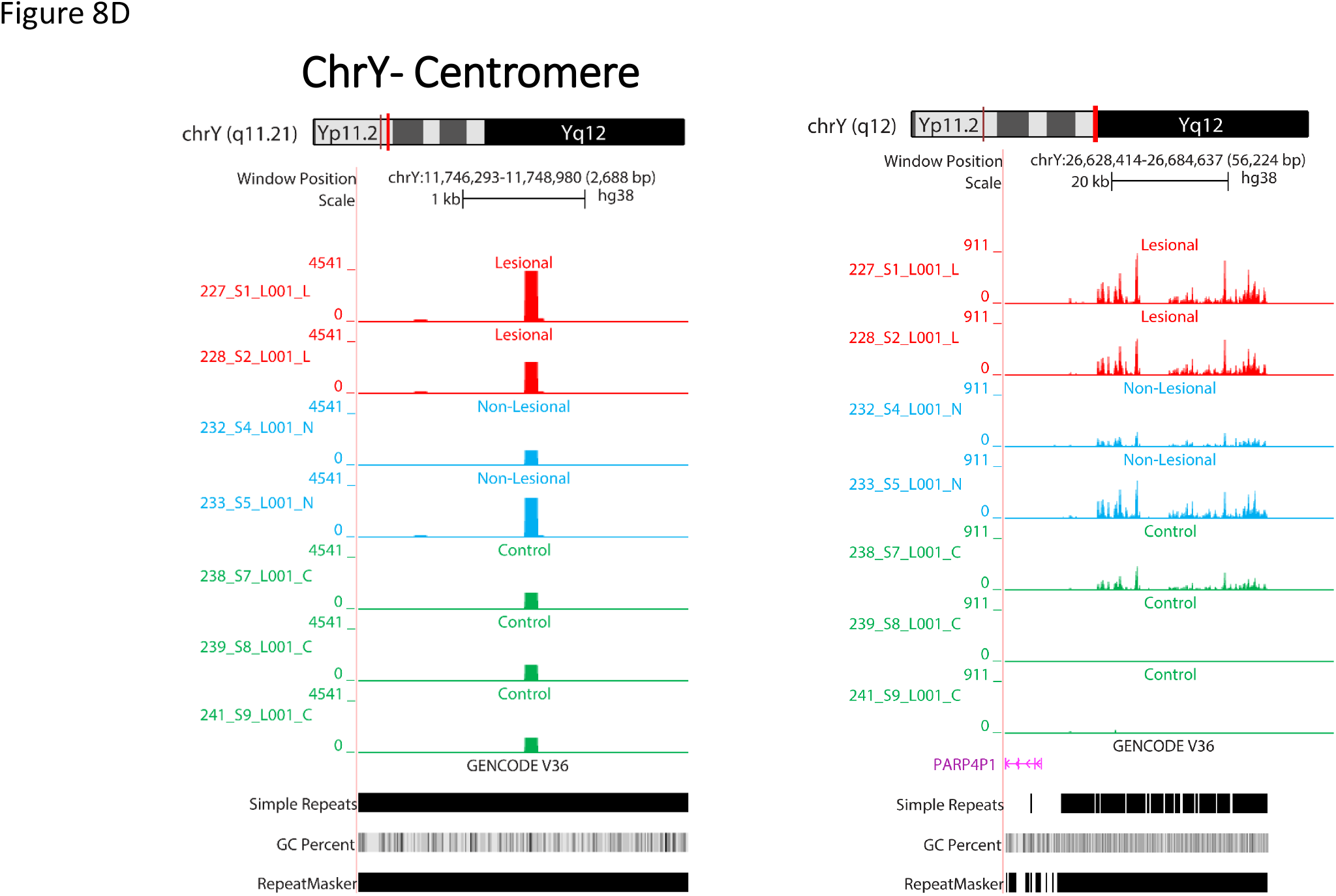
Centromeric and pericentric RNAs profiling genome-wide of DNA-bound RNA molecules. The browser of expression patterns of human DNA associated RNA samples over centromere regions of chromosomes 4, 5, 12, 13, 20, 21, and Y are shown at Figure 8A-D (Sample groups are viewing auto-scaled, each track is auto-scaled to display the track’s highest value). The upper track of each panel shows the chromosome number and the band which centromeric regions are located at based on GRCh38 assembly. The red, blue, and green tracks show expression signals in composite tracks (sample groups auto-scaled). The annotation track is also displayed in pack mode at the bottom of each window (Genecode V36). The upper track of each panel shows the whole chromosome band, which the centromere is depicted in red at the middle of it and the visualized signals are focused on centromeric regions. Simple repeats, GC percent and repeat maskers are also shown at the bottom of the figure.

It is accepted that RNAs are associated with centromeric chromatin. In humans, R-loops are detected at centromeres in mitosis; and an R-loop-driven signaling pathway promotes faithful chromosome segregation and genome stability^39^. It is a common feature that the centromeric regions are desertic in the gene^40,41^. In our data, centromeric RNAs fall in gene-free regions. As shown in Figures 8, RNAs signals are at flanking pericentric heterochromatic of centromeres.

With deep-sequencing analysis, we detect signals at the centromeric regions of chromosomes 4, 5, 12, 13, 20, 21, and Y, of all DNA-bound RNA samples with highly significant enriched repeat sequences of CCATT/GGTAA (see Figure 8A-D). Most of them are located at up-stream regions or promoters so they may have a role in regulating gene expression. Interestingly, centromere read density signals of chromosome 13 are overlapped with the FAM230C lncRNA. On the centromeric region of chromosome 20, two enriched signals were found: one is 79 bp nearby the FRG1Cp pseudogene, and in another one, the repetitive CCATT/GGTAA sequences are FRG1DP (another pseudogene). Centromeric regions sequences of DRNAs are overlapped to simple repeats sequences as shown using the UCSC genome browser.

These observations are in accordance with previous reports that the centromeric sequences have a repetitive nature (alpha-satellite tandem repeats)^42^. The centromeric expression signals of our RNA-seq data overlap with repeat markers including simple repeats (micro-satellites, Figure 8A-D). It has been determined that despite the repetitive sequences, various types of centromeric RNAs play an important role in centromeres as centromeric regions are transcriptionally active, and transcripts are processed, including small RNAs, lncRNAs, circRNAs and hybrids DNA/RNA, which are associated with centromeric proteins and pericentric heterochromatin^42^. It has been shown that the accumulation of R-loops at the centromeric chromatin can alter the integrity of the genome in yeast and mammals^42^. We have also noticed that high transcript signals in the DRNA fraction are visualized in the centromere regions of many chromosomes with higher levels in LA and NL than healthy controls. Our hypothesis was that the accumulation of DNA/RNA hybrids in patients with psoriasis may lead to chromosomal instability, which can result in the development of psoriasis. However, this finding needs to be further investigated by experiments, to have a clue about the molecular function of centromeric R-loops. The idea of the function of centromeres may be an attractive line for future research. When centromeric regions are compared in Figure 8A-D, centromeric RNA signals are generally increased in LA tissue relative to NL and control tissue. In addition, centromeric RNA is relatively high in NL tissue compared to controls.

Evaluation of the skin genome transcripts here by deep sequencing revealed that the amount of non-coding RNAs such as TERRA and centromeres is retained with DNA and is generally increased in psoriasis samples compared to controls (Figure 6 to 8).

### Decreased transcription levels of Ribonuclease HII in patients with psoriasis

Here we report an increase in the level of lncRNAs (telomeres and centromeres regions) associated with DNA, which means an increase in the non-physiological formation of the R-loop that could trigger genome fragility. RAD51 and its partner Breast Cancer 2 (BRCA2) interact with TERRA and promote the association of TERRA with short telomeres^23^. On the other hand, factors in particular, Ribonuclease H and TRF1 oppose the formation of the R-loop^43,20,44,45,46,47^. These proteins play an important role in the physiology of telomeres. We then used the RT-qPCR technique to assess the transcript levels of *Rad51* and of Ribonuclease HII (*RNase HII*) in patients with psoriasis. The results are shown in Figure 9. While the decrease in the level of transcripts in *RNase HII* is highly significant especially in the LA tissues of psoriasis patients, in contrast the levels of Rad51 transcripts do not seem to increase. According to these results, correlation analysis was done and the positive and moderate relation was detected between telomere length and *RNase HII* expression in PSO lesional tissue (p=0.001 rho=0.688). Taken together, these results indicate that an increase in DNA-associated lncRNAs levels could be due to the decreased levels of *RNase HII* transcription in skin tissues of patients with psoriasis. In addition, we determined the level of expression of the *RAD51 and RNase HII* transcripts in blood samples from PSO patients and controls. Patients with PSO have a lower expression level of *RAD51* and *RNase HII* transcripts compared to healthy control; but these results are not significant.

**Figure 9:**
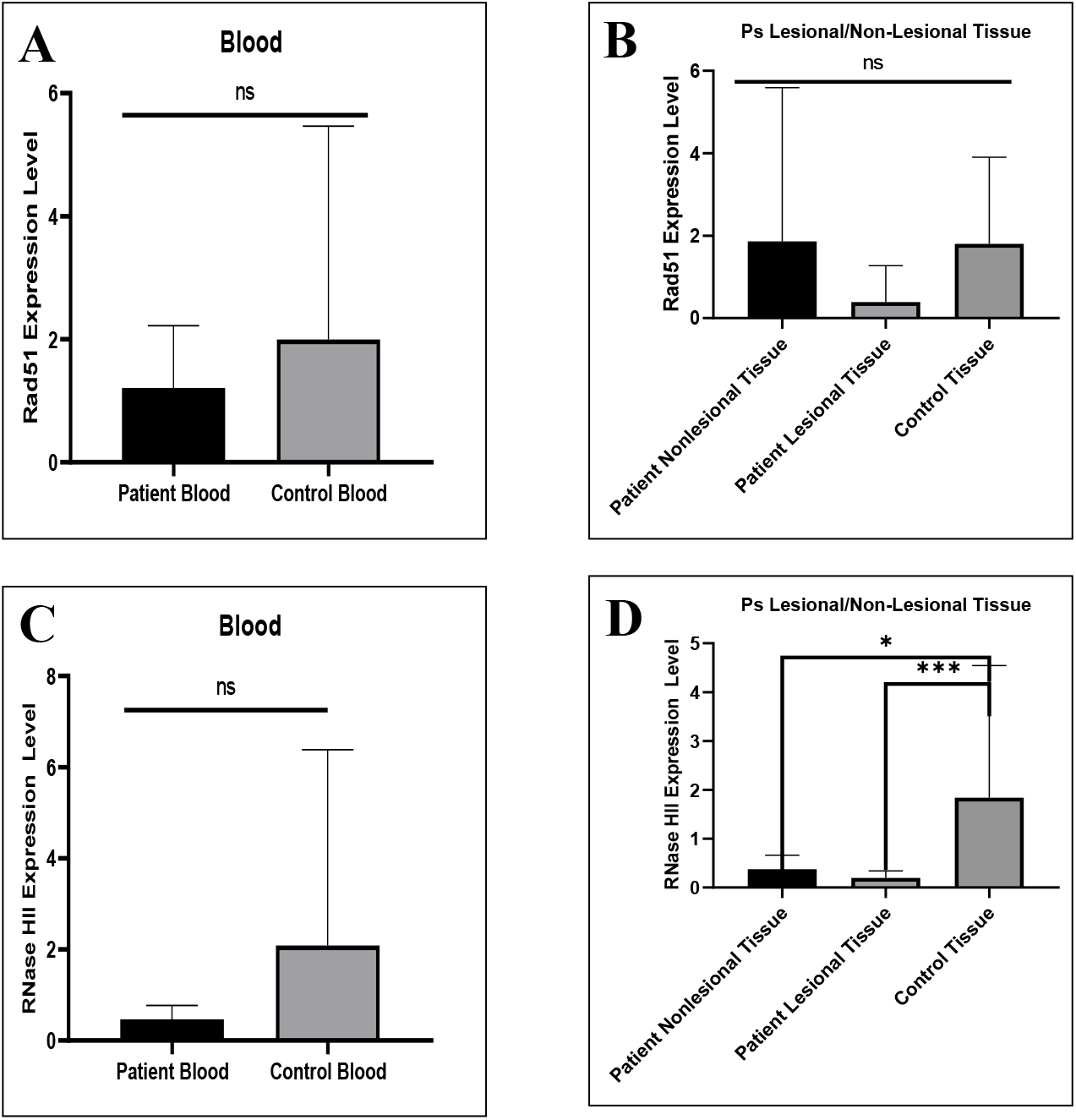
The level of Rad51 mRNA transcript expression in blood samples and lesional/ non lesional tissues. A) The level of Rad51 expression in blood samples compare to healty control (p>0.05) B) The level Rad51 expression in lesional and non lesional tissue compare with healty control (p>0.05). **The level of RNase HII transcript expression in blood samples and lesional/ non lesional tissues. C)** The level RNase HII expression in blood samples compare to healty control (p>0.05) **D)** The level RNase HII expression in lesional and non lesional skin tissue compare to healty control (ps non-lesional vs Control: p=0.04; ps lesional vs Control: p<0.0001).

## Discussion

The aim of this study was to measure the level of TERRA associated with telomeres and others non-coding RNAs (centromeric) to learn more about the state of psoriasis. We hypothesized that psoriasis cells are known to develop short telomeres after lesions contain one or more epigenetic changes that make them susceptible to lesion under certain conditions. Our results indicate that the retention of lncRNAs such as TERRA in telomeres could predict the severity and occurrence of the lesion, or possibly be used as a target in the search for therapeutic or transgenerational follow-ups.

### Telomere length

As it is known, telomeres, which are ribo-nucleoprotein structures at the ends of linear chromosomes, prevent the natural ends of chromosomes from being recognized as DNA breaks and can lead to inappropriate repair activities such as telomere dysfunction, homologous recombination (HR) and non-homologous junction. Due to incomplete DNA replication and nucleolytic degradation, telomeres shorten with each replication cycle, ultimately leading to cell death known as replicative senescence or apoptosis^22^.

In psoriasis, keratinocytes proliferate faster than in normal people. This situation is caused by the formation of keratinized tissue in patients with psoriasis, in fact, due to the increased rate of proliferation and accelerated death of keratinocytes resulting from this rapid cell proliferation, the barrier skin deteriorates, and inflammation occurs in patients.

This indicates that either certain localized factors cause a further shortening of the telomere length in the lesion tissue, or because the keratinocytes in the lesioned area continue to divide. We can only characterize or define this condition by determining the length of the telomere from the biopsy taken from the time where the existing lesion disappears after treatment.

### TERRA profiles

In this study, analysis of TERRA expression profiles of the skin revealed that the increased level of RNA attachment to telomeres correlated with the state of psoriasis. We evidenced that the amount of TERRA associated with DNA was higher in patients with psoriasis even in NL than in healthy controls. Interestingly, the regulation of TERRA levels depends on the location of TERRA in loci associated with chromosomes that exhibit possible genomic alterations, localized epigenetic regulatory changes and TL abnormalities involved in the disease. Genomic alterations cause abnormal TL. In addition, the erosion of telomeres length has a disease promoting function. In particular, damaged skins progress to telomeric shortening. These results supported the hypothesis that increasing the level of TERRA in the R-loop structure compromises the long-term maintenance of TL and cell survival. These skins psoriasis cells do not even –over-express TERRA overall compared to the control, but it may be because of epigenomic changes that they are retained on the telomeres. The shortening of TL in LA skin is evident, but we do not yet know the TL after healing at the lesion points of the skin. TERRA retained in the chromosome is already high in patients with psoriasis, and how TL is controlled in scarred spots requires further study. It has been reported that the lesions promote and accelerate cell proliferation in culture experiments. The downstream signaling pathway of cell growth acceleration has not yet been clarified, but several reports have shown that short telomeres play a role in triggering cell replication^14,15,22^. Here we addressed that higher TERRA retention was generally associated with psoriasis patients suggesting that this could be a potential marker of determinant effects. Indeed, RNA-seq data also confirmed that higher levels of TERRA in the telomeres of patients with psoriasis were associated with increased recurrence. Interestingly, TERRA telomere retention increases in early-stage patients as well as in advanced stage patients. These results are consistent with previous reports by Wu et al,^33^ on telomeres shortening in patient blood cells, and now with increasing TERRA levels could be a coherent epigenetic event for early detection of psoriasis. Thus, studies on TERRA retention on telomeres and others lncRNA on the genome might help to follow-up in trans-generational studies.

We have observed a relationship between TERRA retention and disease progression, but the detailed mechanisms by which skin cells retain TERRA in telomeres remain unknown. Further studies are needed to clarify the relationship between TERRA and intercellular signaling. In addition, TERRAs are not the only lncRNA retained with DNA in patients with psoriasis, in fact, centromeric regions also exhibit higher levels of RNA retention on DNA. This suggests an alteration of the general pathways of resolution of hybrid DNA/RNA structures.

### R-loop structures

Several factors are involved in the regulation of physiological R-loop formation and resolution for review see references included^48^. Our findings indicate decreased levels of important transcripts of Ribonuclease HII (*RNase HII*) enzyme, which removes transcripts from the genome before DNA replication. The reduced levels of *RNase HII* transcripts in the skin of patients with psoriasis help to explain the higher levels of accumulation of lncRNA with DNA. The molecular mechanism of deregulation of *RNase HII* transcripts requires more investigation. Down-regulation at the transcript level could well occur with deregulation of small non-coding RNAs at post-transcriptional levels or with a silencing event at the start of transcription.

Psoriasis is not yet a curable disease, and current treatments block intermediary molecules that play a role in the mechanism of psoriasis or reduce the severity of the disease by suppressing or reducing the proliferation of keratinocytes. Psoriasis seriously and negatively affects the quality of life of patients. Thus, it is clearly very important to develop new strategies for the treatment of this disease.

In conclusion, the levels of TERRA and centromeric regions were higher in RNA attached to DNA from psoriasis patients, regardless of clinical stage thereby making lncRNA as DRNA a potential epigenetic marker and a therapeutic target.

## MATERIALS AND METHODS

Patients diagnosed with psoriasis and healthy volunteers admitted to the Skin and Venereal Dermatology unit of the Faculty of Medicine at Erciyes University were included in this study after approval by the University’s Human Ethics committee: Erciyes University with decision no 17/002.

### Patient Selection

Skin biopsy samples were taken from the area with (LA) and without lesions (NL) from patients with psoriasis, who agreed to participate in the study admitted to the department of Dermatology of the Faculty of Medicine at Erciyes University. The research method of the patients included in the study was prospective, and they had a clinical diagnosis of psoriasis. After calculating the psoriasis area severity index (PASI)^49^ value of patients who did not receive systemic therapy in the past 4 weeks, patients with a PASI value greater than 10 were included in the study.

A total of 35 people over the age of 18 were included in the study, including 20 patients diagnosed with chronic plaque-type psoriasis and 15 healthy volunteers who were consistent with age and gender as a control group. Patients younger than 18 years old, patients with an inflammatory/autoimmune or chronic disease other than psoriasis, and patients who received topical therapy for psoriasis in the last 2 weeks or systemic therapy in the last 4 weeks were not included in the study.

### Taking Skin Biopsy Samples

Skin biopsy samples of LA and NL areas of 20 psoriasis patients included in the study. Tissue samples were taken from the NL skin adjacent to this lesion and from the normal skin area behind the knee (popliteal fossa) of the volunteers in the control group, with a 2 mm punch tool after local anesthesia was applied under the skin, and completely sterile conditions were provided. Tissue samples from patients and controls were immediately delivered to the Genome and Stem Cell Center Genome Unit. Half of the samples were used for DNA isolation and DNA/RNA hybrid determination to assess TL and hybrid TERRA amount, and the other half for RNA isolation to determine the total expression of TERRA. The isolated samples were stored at -80°C.

### DNA and DNA/RNA hybrid isolation

DNA, RNA and DNA/RNA hybrid isolation was performed in accordance with the manufacturer’s protocol using the ZR-Duet DNA/RNA MiniPrep Plus (ZYMO Research (www.zymoresearch.com ZR-Duet ™ DNA/RNA MiniPrep Plus catalog number D7003) Kit from tissue samples. The concentrations and purity of DNA, RNA and DNA/RNA hybrid samples were measured in the Biotech Biospec-Nanodevice.

#### Determination of telomere length

Telomere lengths from obtained DNA samples were determined in a quantitative real-time Light Cycler 480 PCR (Roche, Germany) device using appropriate primers (Tel F:CGGTTTGTTTGGGTTTGGGTTTGGGTTTGGGTTTGGGT and Tel R:GGCTTGCCTTACCCTTACCCTTACCCTTACCCTTACCCT) and standards^50^. Telomere lengths were determined using the house-keeping gene (36B4F: CAGCAAGTGGGAAGGTGTAATCC and 36B4R: CCCATTCTATCATCAACGGGTACA) and then proportioning the obtained Ct values to the 36B4 reference gene.

### Quantitative real-time PCR (RT-qPCR)

In order to determine the TERRA expression level from total RNA and DNA/RNA hybrid samples obtained from the patient and control groups; cDNA synthesis was performed using the EvoScript Reverse Transcriptase cDNA synthesis (07912374001, Roche, Germany) kit. The cDNA samples of the total RNA and DNA/RNA hybrids of the patient and control groups obtained were determined in the Roche LightCycler® 480 Real Time PCR device using previously reported^37^ primers RNase HII (F: GACCCTATTGGAGAGCGAGC and R: TATTTGACCCGCCCAAGCAT), Rad51 (F: GGCCATTAGCCCTTCACCAT and R: TCTGCAAGTGGGACTTTCCT) and GAPDH as the house-keeping gene. All samples were normalized using the data 2^−Δ*Ct*^method after a double study^51^.

### Determination of Genes Expression Levels

RT-qPCR were performed using the high-throughput Light Cycler 480 II Real-Time PCR system (Roche, Germany, Mannheim). In order to determine the genes expression levels of total RNA samples obtained from patient and control groups, cDNA synthesis was performed using the EvoScript Reverse Transcriptase cDNA synthesis (07912374001, Roche, Germany) kit. cDNAs were diluted with nuclease free water in 1:5 ratio. SYBR Green Master (Roche, Germany, Mannheim, Cat No: 04707516001) was used to determine mRNA expression levels of *Rad51* and *RNAse HII* genes in total RNA of tissues and blood samples.

The reaction mix was prepared according to the manufacturer’s instructions: 10 μl of 2x SYBR Green mix, 5 μl of nuclease free water and 0.5 μl of 10 pmol primer assays (*Rad51* F: GGCCATTAGCCCTTCACCAT and Rad 51 R: TCTGCAAGTGGGACTTTCCT; *RNase HII* F: GACCCTATTGGAGAGCGAGC and RNase HII R: TATTTGACCCGCCCAAGCAT) were mixed and dispensed into 96 multi-well plates. The total volume was completed to 20 μl by adding 4 μl of cDNA on it. Thermal cycling conditions consisted of an initial denaturation step at 95°C for 5 min, followed by 40 cycles of 94°C for 20 sec, 60°C for 20 sec, 72°C for 45 sn and 95°C for 15 sec, and finally melting curve was performed at 67°C for 01 sn (melting curve) and cooled at 40°C for 30 sec. Human beta-actin (*ACTB* F: CTCGCCTTTGCCGATCC and ACTB R: TCTCCATGTCGTCCCAGTTG) was used as the reference gene. Changes in gene expression were determined using the 2^-ΔΔCt^ method of relative quantification in all groups.

### Statistical Analysis

The suitability of the data for normal distribution was evaluated by histogram, q-q graphs and Shapiro-Wilk test, and Variance homogeneity with Levene’s test. A two-sample t-test was applied to compare the differences in telomere lengths and TERRA expression of lesional and non-lesional tissues among patients. A One-Way ANOVA test was used to multiply compare Telomere lengths and TERRA expressions between patients and the control group. The relationships between quantitative data were evaluated by Spearman correlation analysis. Data were treated using Graphpad Prism (version 8.0.1) software. P values were considered statistically significant as <0.05. One-way ANOVA analysis of peak annotation data were carried out using the SPSS version 19.0 software. The mean of all the sample groups psoriasis patients and healthy controls data were compared using ANOVA followed by Duncan’s multiple comparison tests. The data were presented as means ± SD.

### RNA library and sequencing

DNA-associated RNA molecules were purified from 9 skin sample biopsies (3 Control (C); 3 lesional (LA); 3 non-lesional (NL). After digestion of DNase, 10-100 ng of RNA were obtained.

Plateform Génomique Institut de Biologie – IBENS (France) performed libraries of all samples corresponding to DNA-associated RNA molecules and small RNA high-throughput sequencing (high-throughput sequencing on Illumina Hiseq 2500 or Illumina MiSeq). Totally, 14 million (m) paired-end reads were mapped to unique sites in the human genome (1L), 20m (2L), 8m (1N), 17m (2N), 7m (2C), 8m (3C), 10m (5C) per sample. Average length of the reads was about 75 nt. All the primary sequence characteristics for sample libraries of the DNA-bound RNA sequences of psoriasis skin samples are summarized in table S1.

### Sequence analysis

FastQC version 0.11.7 was used to do some quality control on the RAW sequencing data. Illumina adapters sequences removed by cutadapt v1.16. Based on the FastQC results, trimming of bad-quality reads was performed using Trimmomatic version 0.36 (10 nucleotides were cropped from the 5’ end of each read, and trimmed bases with Phred score lower than 20 from heading and trailing of each read and the trimmed reads with a size less than 30 bp were removed)^52^. Then aligned to the reference assembly (GRCh38) using Hisat2^53^. Quality of alignment assessed using plot-bamstats utilities of samtools {Li, 2011 #10323}. Bam files converted to BigWig format using deeptools. Visualization of BigWig files was performed using web-based UCSC genome browser (http://genome.ucsc.edu)^54,55^.

### HOMER version 4.9 was used for peak finding

Genome-wide locations of TERRA repeats were found by aligning sequences with lengths of 24 and 48 nucleotides composed of 4 and 8 copies of TERRA to the reference genome using bowtie2 using “-a” argument to report all alignment^31,35^.

Statistically significant peaks of expression were identified using HOMER (10000 size of region used for local filtering, 4-fold over local region, Poisson p-value over local region < 0.0001, false discovery rate (FDR) rate threshold < 0.001).

### Peak finding

To identify significant peaks, we looked for genome-wide enriched peaks. This analysis aimed to efficiently visualize the peaks in terms of genomic region characteristics. The results are consistent between replicates, confirming that the technique is generally very reliable. To find enriched peaks and regions associated with nearby genes and genomic features in the genome using Homer, “findPeaks –o auto” called out was used to perform peak calling and transcript identification analysis. Then to annotating regions in the genome “annotatePeaks.pl” called for performing peak annotation. FDR cutoff of 0.001 used for significant peaks identification. Annotated positions for different regions plotted. The percent of significant peaks located in each genomic region is visualized. Annotated positions for exons, intergenic, intron, promoter-TSS, and TTS are based on the hg38 genomic features Peak annotation.

### Statistical analysis for genomic data

Because of the rheostat-type (non-normal) distributions and the limited number of patients, non-parametric methods were used, and a median value comparison was performed by Wilcoxon rank analysis. Statistical analyses were performed with an unpaired t-test with Welch’s corrections. Data are expressed as the median with p-values <0.05 considered to be statistically significant. The standard error of the mean (SEM) represented as mean±SD.

### Data access

We are waiting for accession number:http://www.ncbi.nlm.nih.gov/geo/query/acc.cgi?acc=GSE166636 On GEO with the GEO accession number GSEXXXX, although it will be private until xxx, however if manuscript is accepted will immediately be made public.

## Data availability

All data are available from the corresponding author upon request and the data related to the RNA-seq experiment are deposited in GEO reference: will be provided soon. Source data underlying the main and supplementary figures are available in Supplementary Data 1.

## Acknowledgments

We are grateful to Yusuf Ozkul for open access to all GenKok Institute Kayseri Turkey. This work is supported by grant 2019-2020 of La Fondation Nestlé France to MR. In addition, this project was supported by TYL-2019-9224, Serpil TAHERİ, Erciyes University Scientific Research Projects Coordination Unit.

## Author contributions statements

Skin Samples provided by MB and SB. Data collection, data analysis and interpretation for molecular studies was performed by EM. Expression analysis for RNase HII and Rad51 was performed by ZY. Figures EM, LK and ZY. Bioinformatic analysis for RNA-Seq were performed by LK. This study conceived and conducted by MR and ST. The project and manuscript were written by MR and ST. All authors reviewed the results and approved the final version of the manuscript.

## “Additional Information”

The experiments were approved by the Turkish Ethics Committee decision number: 17/002 see documents in Supplementary info.

Additional information Supplementary information

The online version contains supplementary material available at:

Correspondence and requests for materials should be addressed to MR

